# Targeting PTRAMP-CSS potently inhibits *P. falciparum* across blood, liver and mosquito stages

**DOI:** 10.64898/2026.05.10.724175

**Authors:** Pailene S. Lim, Nicolai C. Jung, Mikha Gabriela, Jelte M. M. Krol, Kitsanapong Reaksudsan, Myo T. Naung, Danushka S. Marapana, Rainbow W.B. Chan, Diana Nyabundi, Jedida Mwacharo, Melissa Kapulu, Philip Bejon, Francis M. Ndungu, Alyssa E. Barry, Rajagopal Murugan, Annie S. P. Yang, Alan F. Cowman, Stephen W. Scally

## Abstract

Malaria, caused by *Plasmodium falciparum* spans liver, blood, and mosquito stages, limiting the effectiveness of single-stage vaccines. The PTRAMP-CSS heterodimer, a core component of the essential PCRCR invasion complex, is expressed on merozoites, mature gametocytes, and salivary gland sporozoites, enabling single-antigen targeting across multiple lifecycle stages. Nanobodies against PTRAMP-CSS block merozoite invasion of erythrocytes, reduce mosquito infection in membrane-feeding assays, and inhibit sporozoite invasion of primary human hepatocytes. High-resolution crystal structures of inhibitory and non-inhibitory nanobody-antigen complexes identify conserved inhibitory epitopes and guide the design of bispecific nanobody Fc constructs with enhanced potency. In semi-immune Kenyan CHMI samples, higher baseline IgG to PTRAMP-CSS and Ripr is associated with improved parasite control. By demonstrating conserved vulnerability across all three major lifecycle stages, PTRAMP-CSS offers a realistic path to single-antigen, multistage vaccines and biologics that aim to prevent disease and block transmission.

## Introduction

Malaria remains a major global health problem, causing more than 600,000 deaths each year (*1*). The most severe form of malaria is caused by *Plasmodium falciparum*, for which two pre-erythrocytic malaria vaccines, RTS,S/AS01 and R21/Matrix-M^TM^ have received World Health Organisation (WHO) pre-qualification for use in young children within highly malaria endemic regions of Africa (*2*, *3*). Both vaccines target the circumsporozoite protein (CSP) and, while they prevent a high proportion of clinical episodes in seasonal settings, efficacy wanes substantially within a year for RTS,S (*3*, *4*). Furthermore, leading monoclonal antibodies (mAbs) to CSP have demonstrated protective efficacy, providing proof-of-principle for antibody-based prevention (*5*, *6*). Given that sterile immunity or durable protection is hard to achieve by targeting a single stage, antigens that can engage across liver, blood and mosquito stages offer a rare opportunity: a single-antigen strategy that could both reduce disease and block transmission which would materially simplify vaccine or biologic design and increase the potential of a broad and durable impact on malaria elimination.

The PCRCR complex is essential for *P. falciparum* merozoite invasion of erythrocytes and is highly conserved, making it a promising vaccine candidate (*7*, *8*). The complex comprises five proteins, PTRAMP (*Plasmodium* thrombospondin-related apical merozoite protein) (*9*), CSS (cysteine-rich, small, secreted) (*9*), Ripr (Rh5-interacting protein) (*10*), CyRPA (cysteine-rich protective antigen) (*8*, *11*) and Rh5 (reticulocyte-binding protein homologue 5) (*7*, *12*) and binds basigin on the surface of erythrocytes (*13*, *14*). All five proteins are the target of inhibitory antibodies or nanobodies, supporting their potential as vaccine or biologic targets (*9*, *15–19*). Rh5 is the most advanced of these candidates, demonstrating 55% protection against clinical malaria in a Phase IIb clinical trial (*20*). However, as Rh5 is expressed only during the blood stage, Rh5-targeted antibodies cannot act on sporozoites or gametocytes. Consequently, this antigen alone would not provide pre-erythrocytic protection or block transmission, which may limit its overall protective efficacy. In contrast to Rh5, PTRAMP and CSS, which form a disulfide-linked heterodimer, are known to be expressed in merozoites, and whole transcriptome and proteome data suggests they are expressed in late-stage gametocytes and salivary gland sporozoites (*21*, *22*), and therefore may contribute to broader, multi-stage vaccine or biologic strategies.

In this study, we demonstrate that the disulfide-linked PTRAMP-CSS heterodimer is targetable across *P. falciparum* blood, transmission and pre-erythrocytic stages. Nanobodies potently block merozoite erythrocyte invasion in the blood stage, substantially reduce mosquito infection, and inhibit sporozoite invasion into primary human hepatocytes. Seven high-resolution structures delineate conserved inhibitory epitopes and enabled bispecific nanobody-Fc (Nb-Fc) designs with enhanced potency. These findings identify PTRAMP-CSS as a tractable *P. falciparum* antigen across liver, blood and mosquito stages, opening a realistic route to single-antigen, multistage vaccine and biologic strategies.

## Results

### *P. falciparum* PTRAMP-CSS specific nanobodies bind novel epitopes with high affinity

To map inhibitory epitopes on the *P. falciparum* PTRAMP-CSS heterodimer, an alpaca nanobody (Nb) library was generated and subjected to sequential phage-display panning against the PTRAMP-CSS heterodimer and monomeric PTRAMP. This yielded 25 unique Nb sequences that included 19 that recognize CSS, five that bind PTRAMP and one that binds the heterodimer exclusively (fig. S1). Recombinant Nbs were expressed, purified, and characterized by biolayer interferometry (BLI) to determine affinities for monomeric PTRAMP, monomeric CSS and the PTRAMP-CSS heterodimer (table S1, fig. S1). K_D_ values to the heterodimer ranged from 1.9 to 52 nM; most Nbs bound monomeric and heterodimeric antigens with comparable affinities, except for 1F2 which bound only the heterodimer. Domain mapping showed that of the five α-PTRAMP nanobodies, 2D11 and 3F4 target the thrombospondin-type 1 repeat (TSR) and 2C2, 2D12 and 3E2 bind the growth-factor-like domain (GFD) of PTRAMP (table S2, fig. S1A).

The Nb binding epitopes were further investigated using competition binning experiments (Fig. 1A and fig. S2). Nbs were tested in all pairwise combinations to assess their ability to block Ripr binding to PTRAMP-CSS. Previously mapped CSS binders were included for comparison, notably Nbs B1 and B6 that compete with Ripr and Nb D2, which targets an inhibitory epitope on the first domain (D1) of CSS (*9*). Sixteen of the 25 Nbs bound sites that either competed directly with Ripr or competed with a Ripr-competing Nb. Notably, Nb 1G8 binds a distinct Ripr-competing site. Three Nbs competed with the inhibitory Nb D2 and among the five PTRAMP-binding Nbs four distinct non-Ripr-competing epitopes were identified (Fig. 1A and fig. S2B). Overall, these data provide a broad delineation of the PTRAMP-CSS antigenic landscape and revealed distinct epitopes that can be prioritized for structure-guided vaccine or biologic design.

**Figure 1.**
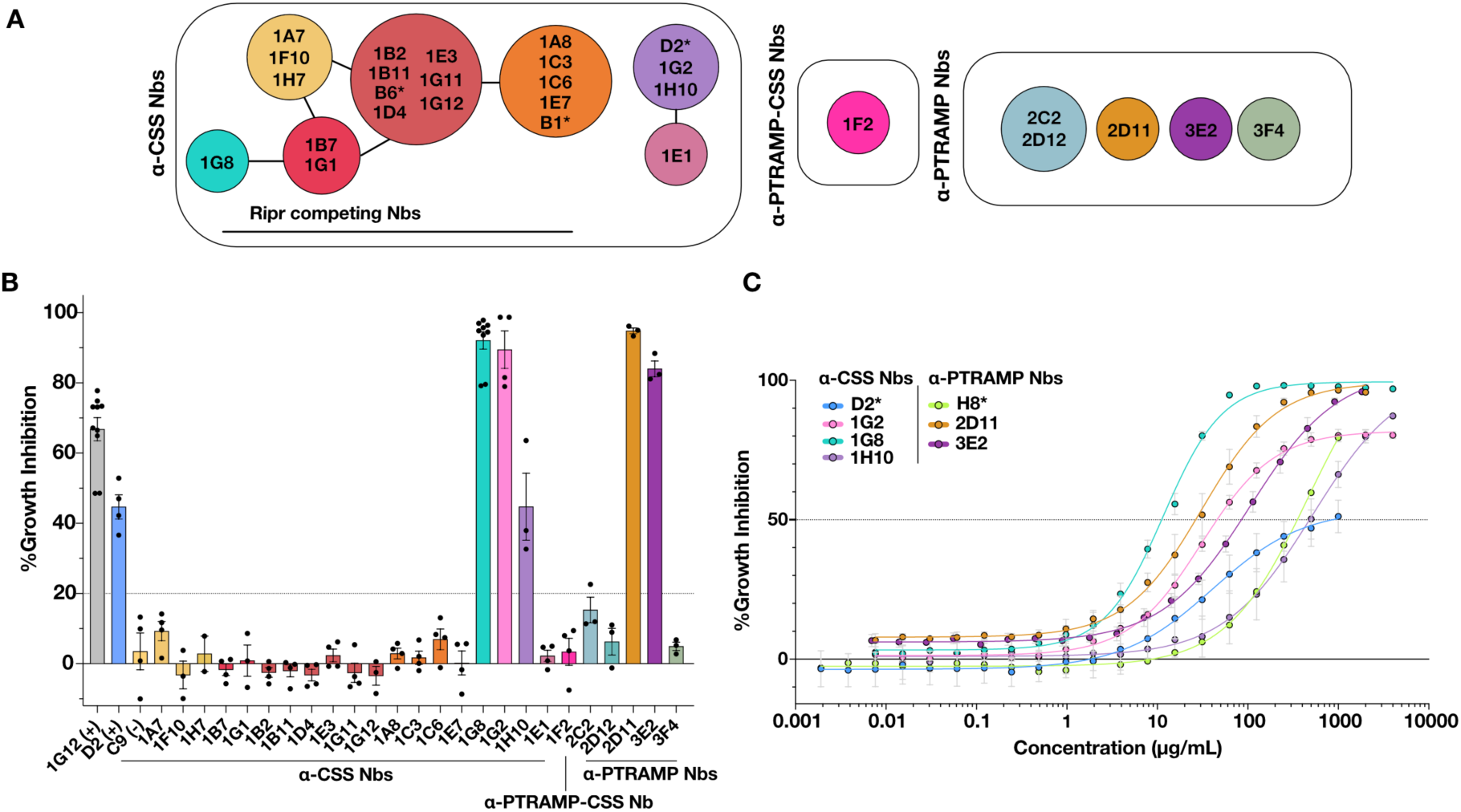
α-PTRAMP-CSS nanobodies potently inhibit blood stage parasite growth in *P. falciparum.* **(A)** Epitope bins of α-PTRAMP and α-CSS Nbs. Each bin is represented by a unique color and competition between Nbs is represented by a black line connecting circles. Nbs that compete with PTRAMP-CSS binding to Ripr are also indicated. Asterisks denote Nbs that were previously characterized (*9*) See also fig. S2. (**B)** Growth inhibition of α-PTRAMP-CSS nanobodies at 0.5 mg/mL. Ripr mAb 1G12 and α-CSS D2 Nb was used as a positive control at 1 mg/mL, whilst α-CSS C9 was used as a negative control at 0.5 mg/mL (*9*, *18*). **(C)** Growth inhibition dilution series for α-PTRAMP Nbs 2D11 and 3E2 and α-CSS nanobodies 1G2, 1G8 and 1H10. Positive controls α-PTRAMP H8 and α-PfCSS D2 nanobodies were included to compare EC_50_ (µg/mL) values (see also table S3).

### PTRAMP-CSS nanobodies potently inhibit blood stage parasite growth

A subset of Nbs showed strong activity in *P. falciparum* growth inhibition assays with five out of 25 reducing parasite growth, comprising three α-CSS Nbs (1G2, 1G8, 1H10) and two α-PTRAMP Nbs (2D11, 3E2) (Fig. 1B). At a single test concentration, 1G2, 1G8, 2D11 and 3E2 each produced >84% inhibition. Notably, this included the Ripr-competing Nb 1G8 suggesting that although almost all Ripr-competing Nbs were non-inhibitory, subtle differences in how they engage the complex may give rise to distinct inhibitory effects.

Inhibitory Nbs against PTRAMP and CSS showed a dose-dependent inhibition when titrated (Fig. 1C). All Nbs except for 1H10 demonstrated improved efficacy from the previously identified inhibitory Nbs D2 and H8 (Fig. 1C and table S3) (*9*). 1G8 was the best performing Nb overall with an EC_50_ of 11.8 µg/mL (95% CI 10.5 - 13.2 µg/mL), a 36-fold improvement compared to the anti-CSS Nb D2 (Fig. 1C and table S3). 1G2 showed a 7-fold improvement in EC_50_ (58.1 µg/mL (95% CI 46.4 - 72.5 µg/mL). 2D11 was the most potent α-PTRAMP Nb with an EC_50_ of 31.4 µg/mL (95% CI 26.9 - 36.6 µg/mL) and a 10-fold improvement in efficacy compared to the anti-PTRAMP Nb H8. Together, these data identify several high-affinity and neutralizing Nbs that target novel epitopes on the PTRAMP-CSS complex and provide validated molecular probes for vaccine design and biologic development.

### High resolution structures define conserved, inhibitory epitopes on the PTRAMP-CSS heterodimer

To define the molecular basis of inhibition, we determined the crystal structures of seven Nbs, 1G2, 1G8, 1H10, 2C2, 2D11, 3E2 and 3F4, in complex with PTRAMP, CSS or the PTRAMP-CSS heterodimer (table S4-S13). Specifically, we determined the structures of anti-PTRAMP Nbs, 2C2, 3E2, 2D11 and 3F4 in complex in full-length PTRAMP (2C2-PTRAMP), GFD (3E2-GFD) and TSR (2D11-3F4-TSR) to resolutions of 1.50 Å, 2.40 Å and 2.12 Å, respectively (Fig. 2A-C and table S4). We determined the structures of anti-CSS Nbs 1G2, 1H10 in complex with CSS to resolutions of 3.23 Å and 2.64 Å, respectively (Figure 2D and E, table S4) and 1G8 in complex with PTRAMP-CSS to a resolution of 4.20 Å (Figure 2F, table S4). Collectively, these crystal structures provide a broad molecular understanding of inhibitory and non-inhibitory epitope landscape on the surface of this essential complex and reveal conserved epitopes than can be directly exploited for vaccine design.

**Figure 2.**
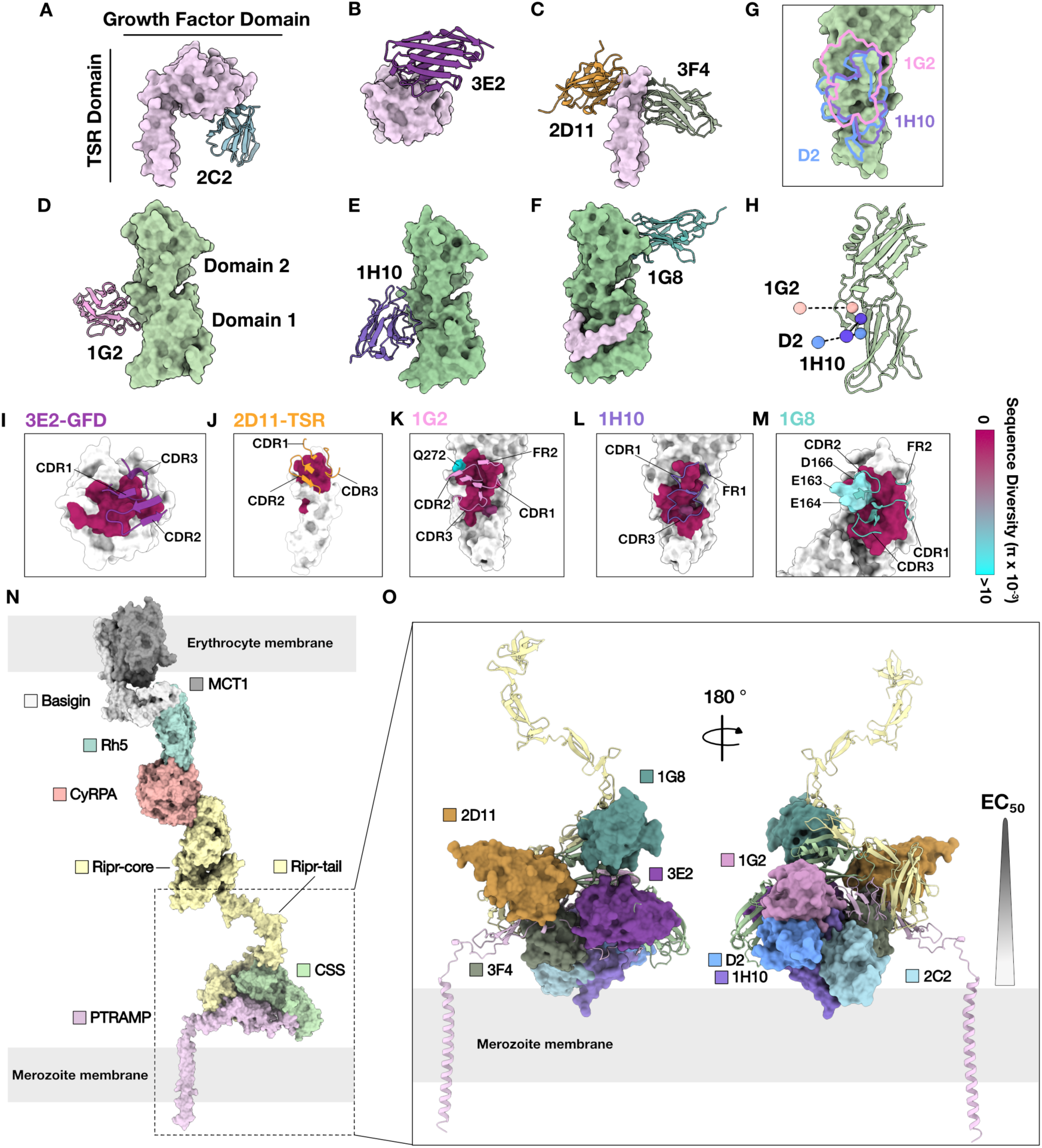
Crystal structures of non-inhibitory and inhibitory PTRAMP-CSS nanobodies in complex with PTRAMP or CSS reveal conserved strain-transcending inhibitory epitopes. **(A)** Crystal structure of 2C2 nanobody with full-length PTRAMP. Growth factor domain (GFD) and thrombospondin type-1 repeat (TSR) domain of PTRAMP are indicated. **(B)** Crystal structures of 3E2 nanobody with PTRAMP GFD and **(C)** 2D11 and 3F4 nanobodies co-complexed with TSR domain. **(D)** Crystal structure of 1G2 nanobody with full-length CSS. **(E)** Crystal structure of 1H10 binding to CSS. **(F)** Crystal structure of 1G8 nanobody with PTRAMP-CSS. **(G)** Superimposed recognition sites of 1G2, 1H10, and D2 nanobodies on CSS. **(H)** Angle of approach of 1G2, 1H10 and D2 Nb binding to CSS. **(I – M)** Interaction of inhibitory Nb CDRs and framework regions (FR) with residues on PTRAMP and CSS (See also fig. S3C and F). Epitopes on PTRAMP or CSS are colored by global sequence diversity, where maroon is highly conserved and cyan is highly diverse (see also fig. S4 and table S14). **(N)** Model of the PCRCR complex binding to basigin on the erythrocyte membrane. The model was generated using structural information from PDB entries 7CKR (MCT1–Basigin) (*71*), 4U0Q (Basigin–Rh5) (*14*), 8CDD (Rh5–CyRPA–Ripr) (*72*), together with AF3 predictions (*23*). **(O)** Mapping of non-inhibitory and inhibitory nanobodies onto the AF3 predicted PTRAMP-CSS-Ripr complex during erythrocyte invasion.

### Crystal structures reveal distinct inhibitory epitopes across the growth-factor and TSR domains of PTRAMP

The 2C2-PTRAMP co-crystal defined the high-resolution architecture of the PTRAMP ectodomain and revealed two discrete subdomains, in agreement with sequence and AlphaFold predictions (*23*). The N-terminal domain exhibits structural similarity to the pro-domain of the transforming growth factor (TGF)-β superfamily, including TGF-β and bone morphogenetic protein 9 (BMP9) (fig. S3A), as well as the carbohydrate binding module (CBM) of carbohydrate binding proteins such as the *Defluviitalea phaphyphila* alignate lyase, Dp0100 (*24*). This is despite a low sequence identity of 11% and 9%, respectively (fig. S3A). The second domain shares structural similarity with TSRs, including human TSR (rmsd ∼2.2 Å), a common structural element seen in many *Plasmodium* species proteins (fig. S3B). Notably, the PTRAMP-TSR domain possesses the canonical WxxWxxC motif.

Consistent with kinetics analyses, 2C2 binds exclusively to the PTRAMP-GFD (Fig. 2A and fig. S1A). Specifically, 2C2 contacts one face of the β-sandwich, including β1’, β2, β3, β5, β6, β8 strands and surrounding loops. Indeed, 2C2 engages with PTRAMP through its FR1, CDR1, CDR2, and CDR3 loops, forming an extensive network of 25 hydrogen bonds and 11 salt bridges, with a buried surface area (BSA) of 889 Å^2^ (fig. S3C and table S5 and S13). Similar to 2C2, 3E2 also binds exclusively to the PTRAMP-GFD (Fig. 2B). However, it primarily binds to loops at the opposite end of the GFD, specifically a short α-helix within the β3-β4 loop, the β7- β8 loop, and residues in the β1’ and β1 strands. 3E2 interacts with the GFD through its CDR1, CDR2, and CDR3 loops (Fig. 2I, fig. S3C and table S6). Despite having nearly half the BSA of 2C2 (461.7 Å^2^), 3E2 is inhibitory while 2C2 was not, suggesting the specific epitope plays a more critical role than BSA in determining the inhibitory effect.

The co-complex of the inhibitory 2D11 and the non-inhibitory 3F4 nanobodies with the TSR domain of PTRAMP bound opposite faces of the structural motif (Fig. 2C). Specifically, 2D11 interacts almost exclusively with the β2- β3 loop through its CDR1, CDR2 and CDR3 loops (BSA of 494.9 Å^2^), which was stabilized by a conserved disulfide bond (Fig. 2J, fig. S3C and table S7). In contrast, 3F4 bound to the first Trp of the WxxWxxC motif of the TSR domain, Trp248 and the interlocking Arg269 from the β2 strand (fig. S3D, E and table S8). Unexpectedly 2D11 and 3F4 interact with each other through heterotypic interactions, involving 65.5 Å^2^ of BSA and two H-bonds mediated by the 2D11 CDR2 and 3F4 CDR3 loops (fig. S3D and table S9). These Nb-PTRAMP structures define multiple, spatially distinct epitopes across the GFD and TSR domains and reveal that inhibitory activity maps to specific, surface-exposed sites rather than to overall binding footprint. These results both inform epitope-focused immunogen design and provide mechanistic probes to dissect the role of PTRAMP in invasion.

### Structural mapping of CSS nanobody complexes identifies a novel, potent domain 2 inhibitory epitope

Crystal structures of 1G2 and 1H10 in complex with CSS revealed that, consistent with the competition binning data, the Nbs bind to an overlapping epitope with D2 (Fig. 2D-G). 1G2 and 1H10 contact both the D1 and D2 degenerate 6-Cys domains of CSS. Specifically, one face of the β-sheet of the D1 domain and a loop from the D2 domain. Indeed, the 1H10 epitope was highly similar to that of D2, with near-complete overlap (Fig. 2G). 1H10 interacts via its CDR1, CDR3 and FR1 loops and a BSA of 733 Å^2^ (Fig 2L, fig. S3F and table S10). In contrast, 1G2 engages more extensively with the D2 domain of CSS, via all CDRs and FR2, with a BSA of 708 Å^2^ (Fig. 2K, fig. S3F and table S11). While all three Nbs target a similar epitope, their angles of approach differ, with 1G2 and D2 adopting a comparable binding approach, whereas 1H10 binds from a distinctly different angle, which may account for its reduced potency (Fig. 2H).

The crystal structure of 1G8 in complex with the PTRAMP-CSS complex revealed that only 1G8, CSS, and 20 amino acids of PTRAMP were resolved in the crystal lattice, corresponding to the region surrounding the disulfide linkage between PTRAMP and CSS (Fig. 2F). In this configuration, PTRAMP adopts an antiparallel β-strand, likely reflecting flexibility in the linker connecting this segment to the core region that interacts with CSS C30 and PTRAMP C60, and consistent with previous observations of the *P. vivax* PTRAMP-CSS heterodimer (*25*). In contrast to 1G2 and 1H10, 1G8 binds to a distal site on CSS, consistent with the competition data. 1G8 binds exclusively to one face of the β-sheet on the D2 domain of CSS, opposite to the 1G2 binding face on D1. 1G8 interacts via all its CDR loops and FR2 region with a BSA of 817 Å^2^ (Fig. 2M, fig. S3F and table S12). The structure of 1G8-CSS has revealed a novel and potent growth inhibitory epitope on the D2 domain of CSS.

### PTRAMP and CSS are highly conserved across global *P. falciparum* strains

To evaluate antigenic conservation, 20,646 *P. falciparum* field isolate genomes were analysed from 24 countries to map observed sequence variation onto the PTRAMP-CSS structures (fig. S4 and table S14) (*26*). In this dataset PTRAMP showed no amino acid polymorphisms, indicating high conservation of the sequence among the natural isolates examined (fig. S4A). In contrast, CSS carried 41 amino acid substitutions in total, but only four variants occurred at appreciable global frequencies: Q272K (67%), K93N (46%), D166E (9.5%) and E163K (8.0%) (fig. S5B and table S14). Q272K and K93N localize to the D1 domain of CSS, whereas D166E and E163K lie on domain 2 (fig. S4B).

Mapping these polymorphisms onto the Nb-PTRAMP-CSS structures indicated that the most potent inhibitory epitopes defined are largely conserved. PTRAMP inhibitory epitopes targeted by α-PTRAMP Nbs have no polymorphisms in this global dataset (Fig. 2I–J and fig. S4A). Of the CSS Nbs, 1H10 engages an epitope that was effectively conserved across the analysed genomes (Fig. 2L and fig. S4B), whereas 1G2 contacts residue Gln272 which is adjacent, but not central to, the main binding interface (Fig. 2K and fig. S4 and 5A). The 1G8 epitope on D2 overlaps a loop spanning residues F161–D166 that shows modest variation; among these positions D166E and E163K are the most frequent (9.5% and 8.0%, respectively) (Fig. 2M, fig. S5B and table S14).

To determine if common polymorphisms would affect 1G2 or 1G8 binding, six PfCSS haplotypes were expressed which collectively contained all the possible mutations identified within the 1G2 and 1G8 binding sites (fig. S5C and D and table S14). No differences in K_D_ values were seen for 1G2 binding to CSS-G224 (containing Q272K mutation) compared to CSS-3D7 (fig. S5E and G, table S15). In contrast, a 4.3 to 37-fold decrease in affinity was seen for CSS variants binding to 1G8 compared to 3D7, with CSS-Guinea (containing E164K and D166E mutations) having the lowest affinity with a K_D_ value of 124 ± 41 nM (fig. S5F and G and table S15). Despite a reduction in binding affinity for 1G8, both 1G8 and 1G2 can still bind strongly to recombinant CSS variants. Whether these mutations affect functional activity of these nanobodies has yet to be determined.

Taken together, these data indicate that PTRAMP is conserved across all analyzed genomes and that CSS is predominantly conserved with a limited number of relatively common, geographically restricted variants. This overall conservation, and the fact that the principal inhibitory epitopes are either conserved or engage residues only adjacent to polymorphic sites, supports the candidacy of PTRAMP-CSS for broadly effective, structure-guided vaccine and biologic design.

### Epitope accessibility and membrane proximity govern nanobody inhibitory activity

To understand how Nbs block erythrocyte invasion by *P. falciparum* merozoites, inhibitory potency (EC_50_) was tested for correlation with kinetic or structural parameters including K_on_, K_off_, K_D_ and BSA. No significant correlations were observed (fig. S6A), indicating that affinity or footprint size alone did not explain functional potency.

To understand how Nbs interact with the PTRAMP-CSS complex in the context of its predicted interaction with Ripr, Nb binding was modelled onto the PTRAMP-CSS-Ripr complex using the AlphaFold3 structural prediction (Fig. 2N and O) (*23*) . Consistent with the binning data, 1G8, the most potent Nb, binds immediately adjacent to the predicted Ripr binding site, and would sterically interfere with the flexible Ripr-tail. The fact that 1G8 remains highly potent in growth assays, suggests it can access the epitope despite this overlap, likely due to local structural flexibility. Indeed, while the PTRAMP-CSS assembly was probably flexible, a striking trend was evident in which Nbs, that bind epitopes farther from the merozoite plasma membrane, tended to be more inhibitory (Fig. 2O and fig. S6B). Non-inhibitory Nbs, 2C2 and 3F4, bind closest to the membrane, followed by intermediary Nbs, including 1H10, D2 and 1G2, while the most inhibitory Nbs, 2D11, 3E2, and 1G8 bind distally (Fig. 2O and fig. S6B). Together, these findings indicate that both epitope accessibility and distance from the merozoite plasma membrane critically determine Nb inhibitory potency.

### Bi-specific Nb-Fc fusion proteins improve the potency in blood stage

Pairwise analyses were performed to assess synergy among inhibitory α-PTRAMP-CSS Nbs, using the Bliss definition of additivity (*15*, *27*). All inhibitory Nbs excluding 1H10 were measured in pair-wise combinations. All Nb combinations did not show a synergistic effect, instead demonstrating Bliss additivity (Fig. 3A and fig. S7). No antagonistic effect was seen which was expected given that these Nbs bind to different epitopes on PTRAMP-CSS and would not compete with one another.

**Figure 3.**
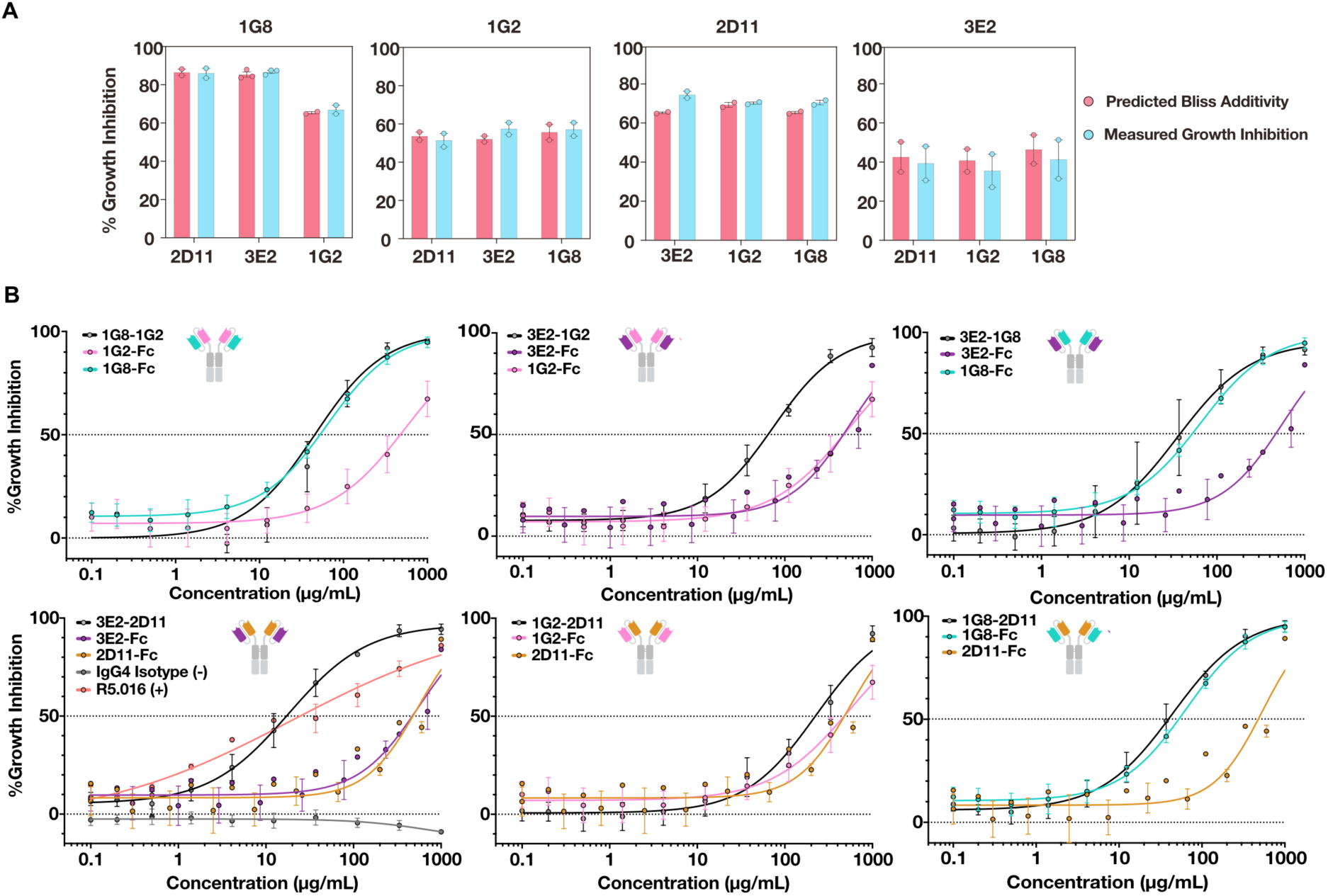
Bispecific Nb-Fc proteins improve the potency of Nbs against *P. falciparum.* **(A)** Predicted GIA based on Bliss Additivity (red) compared to measured GIA (blue) for Nb combinations where one was held at 31.25 µg/mL (title) and the other held at ∼20% GIA (bottom X-axis). Each data point is from one independent experiment performed in triplicate (n = 2). (See also fig. S7.) **(B)** Growth inhibition dilution series comparing individual nanobody-Fc (Nb-Fc) inhibition with bispecific nanobody-Fc (BsNb-Fc) combinations. BsNb-Fcs are shown in black whilst Nb-Fc are colored by epitope bins (Fig. 1A). Anti-PfRh5 mAb R5.011 was included as a positive control (red), and an IgG4 human isotype antibody was used as a negative control (grey). Data shown is the average of three independent experiments performed in triplicate. Error bars show SEM. Four parametric non-linear regression is shown in solid-colored lines.

To evaluate the potential of α-PTRAMP-CSS Nbs as biologics for malaria prevention, the four most potent candidates, 1G2, 1G8, 2D11 and 3E2 were reformatted into bispecific Nb-Fc constructs. For comparison the individual Nb-Fc fusion proteins and R5.016 were tested, one of the most potent Rh5 human mAb described to date (*15*). Affinity studies comparing Nb-Fc and BsNb-Fc binding to PTRAMP-CSS demonstrate a 5 to 7,500-fold improvement in K_D_, likely due to increased avidity (fig. S8 and table S16).

When tested in GIA, two of the six constructs, the 3E2-1G2 and 3E2-2D11, displayed significantly improved EC_50_ values compared to their monospecific counterparts (Fig. 3B and table S17). Notably, the bivalent 3E2-2D11, which contains two α-PTRAMP nanobodies, achieved an EC_50_ of 18.1 µg/mL (162 nM, 95% CI 11.7 – 27.5 µg/mL) (Fig. 3B), representing a 22-fold improvement compared to the individual components, and equivalent to the potency of α-Rh5 mAb R5.016 (EC_50_ 20.7 µg/mL (137 nM), EC_50_ 9 µg/mL (60 nM) reported previously) (*15*). Importantly, 3E2-2D11 exhibited a lower EC_80_ value compared to R5.016 (100 µg/mL (902 nM) vs 400 µg/mL (2.6 µM)), due to a steeper Hill slope, which may prove a more useful measure of efficacy compared to EC_50_ (*15*). Meanwhile, 3E2-1G2, which targets both PTRAMP and CSS, achieved an EC_50_ of 65 µg/mL, representing a 7-fold improvement compared to the individual components. These findings demonstrated that bispecific Nb-Fc constructs substantially enhance potency and broaden inhibitory capacity of anti-PTRAMP-CSS biologics.

### PTRAMP-CSS is expressed in late-stage gametocytes and sporozoites

To assess the feasibility of targeting PTRAMP-CSS in other lifecycle stages and thereby evaluate its potential as a multi-stage malaria vaccine candidate or biologic target, the expression and function of the PTRAMP-CSS complex across transmission stages and pre-erythrocytic stages of the parasite lifecycle was examined. Across gametocyte development (stage II-V), CyRPA and Rh5 transcripts declined, Ripr showed a modest increase, while PTRAMP and CSS transcripts increased (Fig. 4A). Furthermore, immunoblotting with both inhibitory Nbs 1G8, 1G2 and 2D11, and non-inhibitory Nbs 1G12 and 1F2, detected a band corresponding to PTRAMP-CSS, albeit at lower intensity than in schizonts, confirming the Nbs recognize PTRAMP-CSS present in stage V gametocyte as well as in schizont lysates (Fig. 4B). Finally, to assess PTRAMP and CSS expression in sporozoites, salivary gland sporozoites were probed with anti-PTRAMP and anti-CSS mAbs and detected a band corresponding to monomeric PTRAMP and CSS under reducing conditions (Fig. 4C). Collectively, these results demonstrate that PTRAMP and CSS are expressed across all major *P. falciparum* lifecycle stages, supporting their potential as multi-stage targets for malaria intervention.

**Figure 4.**
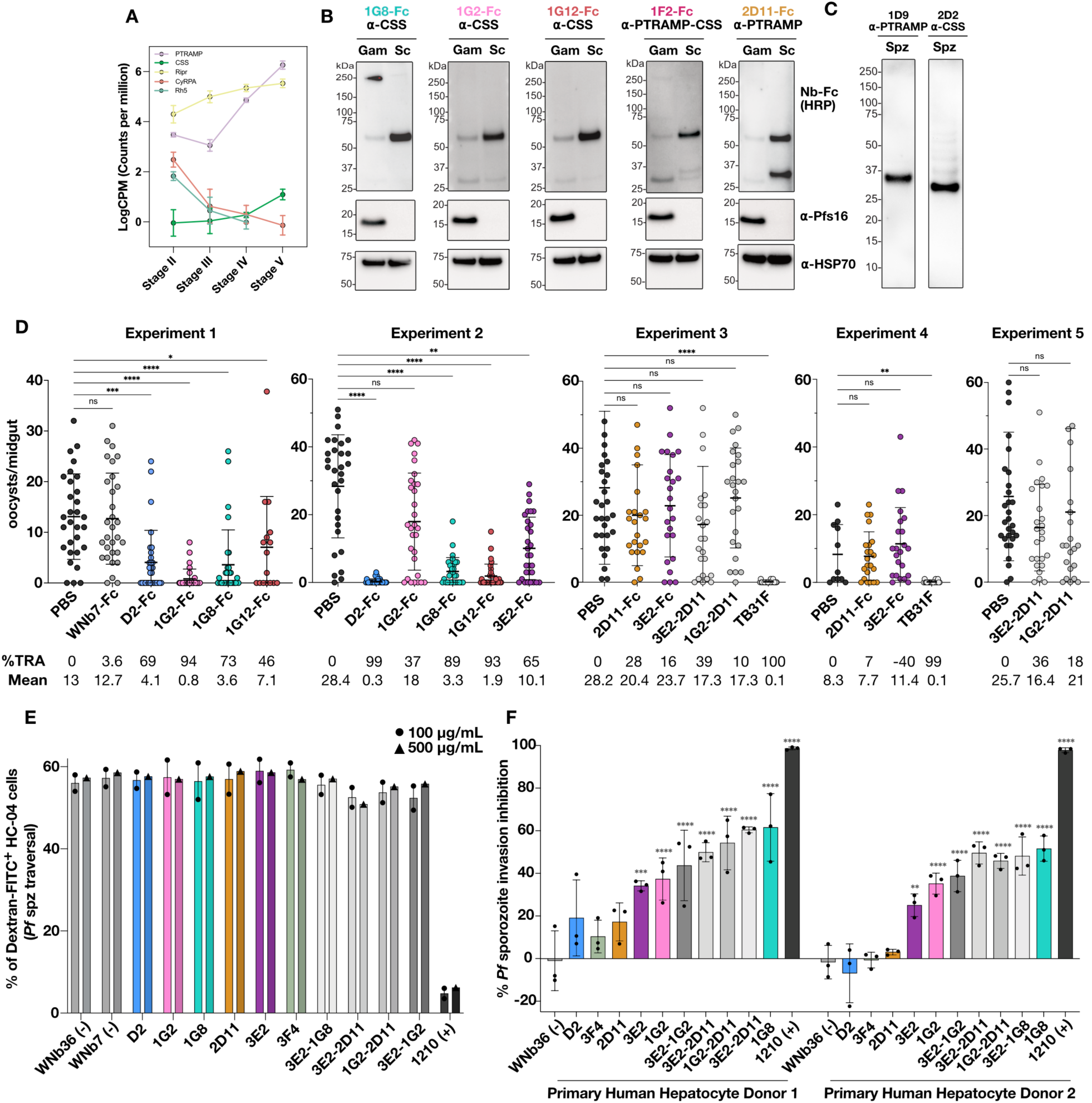
α-PTRAMP-CSS Nb-Fc potently inhibit *P. falciparum* transmission and sporozoite invasion. **(A)** Expression of mRNA transcripts encoding the PTRAMP-CSS-Ripr-Rh5 (PCRCR) protein complex across stage II to V gametocytes (n=5). Data shown is the average log counts per million of each protein across five independent experiments, with error bars representing SEM. **(B)** Western blots of Nb-Fcs against PTRAMP-CSS from *P. falciparum* stage V gametocytes and schizonts lysate (non-reduced) separated by SDS-PAGE, detected with HRP-conjugated anti-human IgG (n = 2). α-Pfs16 Ab was used as a gametocyte marker and α-HSP70 Ab as a loading control. Molecular weight in kDa is shown. **(C)** Western blots of α-PTRAMP mAb 1D9 and α-CSS mAb 2D2 against lysate from 200,000 sporozoites (reduced) separated by SDS-PAGE, detected with goat α-mouse-HRP-conjugated IgG. **(D)** Oocyst counts from five independent standard membrane feeding assays (SMFA) experiments testing α-CSS Nb-Fcs (Experiment 1 and 2) and α-PfPTRAMP Nb-Fcs (Experiments 2, 3, 4 and 5). Mosquitoes were dissected 7 – 8 days post feeding with Stage V NF54 iGP2 gametocytes. α-SARS-CoV-2 WNb7-Fc and α-Pfs25 mAb TB31F were included as negative and positive controls, respectively. Transmission reducing activity and mean number of oocysts per condition and standard deviation are also shown (see also table S18). Statistically significant differences between untreated control (PBS) and treatment with Nb-Fcs was determined by Kruskal-Wallis unpaired test followed by Dunn’s post hoc test. **(E)** Percent dextran positive HC-04.J7 hepatocyte cells indicating *P. falciparum* sporozoite traversal upon treatment with PTRAMP-CSS specific Nb-Fcs and BsNb-Fcs tested at 100 µg/mL (circle) (n = 2) and 500 µg/mL (triangle) (n = 1). All experiments were performed in triplicate. **(F)** Inhibition of sporozoite invasion into hepatocytes using primary human hepatocyte donors (n=2), treated with Nb-Fcs and BsNb-Fcs. Antibody 1210 anti-PfCSP was used as a positive control for both experiments, and WNb7-Fc and WNb36-Fc were used as negative controls. Error bars represent standard deviation. For all analysis of significance, *p*-value < 0.0001 (****), <0.0002 (***), <0.0021 (**), <0.0332 (*), <0.1234 (ns).

### PTRAMP-CSS Nb-Fc fusion proteins potently inhibit transmission of *P. falciparum* in mosquitoes

The transmission blocking capacity of the PTRAMP-CSS specific Nb-Fcs were tested in standard membrane feeding assays (SMFAs). PBS and WNb7-Fc served as negative controls, while TB31F was included as a positive control (*28*, *29*). α-CSS Nb-Fcs D2-Fc, 1G8-Fc and 1G12-Fc demonstrated consistent transmission reducing activity (TRA) that was statistically significant compared with PBS controls. D2-Fc showed a TRA of 84 ± 15%, 1G8-Fc had 81 ± 7.9% TRA, and 1G12-Fc was 70 ± 23% TRA from two independent experiments (Fig. 4D and table S18). Interestingly, although 1G12-Fc was non-inhibitory in blood stage GIAs, it exhibited moderate TRA in SMFAs, suggesting the Ripr binding site on PTRAMP-CSS may be exposed in mosquito-stage parasites and that distinct epitopes are targetable across different lifecycle stages. In contrast, α-PTRAMP Nb-Fcs 3E2 and 2D11 showed moderate to weak TRA (Fig. 4D and table S18). Nevertheless, when combined as a bispecific Nb-Fc, 3E2-2D11-Fc showed moderate TRA of 38 ± 1.3% (n=2), suggesting it was targetable (Fig. 4D and table S18). Overall, the SMFA data suggested the PTRAMP-CSS complex contributes to parasite (oocyst) development in the mosquito, with anti-CSS nanobodies exhibiting greater transmission-blocking potency than anti-PTRAMP nanobodies.

### PTRAMP-CSS Nb-Fc inhibit sporozoite invasion into primary human hepatocytes

To assess whether the PTRAMP-CSS complex contributes to parasite survival during pre-erythrocytic stages, a subset of our Nb-Fcs and BsNb-Fc constructs in functional sporozoite assays were tested. WNb7-Fc and WNb36-Fc served as negative controls, while anti-CSP mAb 1210 was included as a positive control (*28*, *30*). None of the Nb-Fcs or BsNb-Fcs inhibited *P. falciparum* sporozoite traversal through HC-04.J7 hepatocyte cells at either low, 100 µg/mL (n=2), or high 500 µg/mL (n=1) concentrations (Fig. 4E and table S19). In contrast, Nb-Fc 1G8, the most potent blood- and mosquito-stage inhibitor, was the most effective when tested for invasion of primary human hepatocytes, demonstrating 57% inhibition of sporozoite invasion (Fig. 4F and table S20). All BsNb-Fcs constructs tested showed >50 % inhibition of sporozoite invasion. Both 1G2-2D11 and 3E2-2D11 outperformed their monospecific counterparts, highlighting the utility of BsNb-Fcs in inhibiting sporozoite development. In contrast, three Nb-Fcs, D2 (α-CSS), 3F4 and 2D11 (α-PTRAMP) did not show statistically significant inhibition relative to the control (Fig. 4F and table S20).

Overall, these data show that PTRAMP and CSS are expressed and can be inhibited in *P. falciparum* schizonts, mature gametocytes and sporozoites. Importantly, this work provides the first demonstration the highly conserved PTRAMP-CSS complex can be targeted across blood, liver and mosquito stages. The 1G8-Fc construct consistently inhibited all three stages, whereas 1G2-Fc displayed strong blood-stage activity and moderate inhibition of transmission and sporozoite invasion.

### Human IgG antibodies to Ripr and PTRAMP-CSS are associated with better clinical outcomes after sporozoite challenge

To determine whether the PCRCR complex is immunogenic in natural infection and whether specific IgG antibodies are associated with protection, we measured IgG antibodies against recombinant PTRAMP-CSS, Ripr, CyRPA and Rh5 in 137 semi-immune Kenyan adults undergoing controlled human malaria infection (CHMI-SIKA) (*31*). Antibodies were measured at baseline (C-1) and on days 14 and 35 post-challenge. Baseline (C-1) and day-14 IgG levels against Ripr and PTRAMP-CSS were significantly higher in participants who controlled infection below the clinical protocol treatment threshold of 500 parasites/µL (untreated; n = 83) than in those who failed to control parasite growth and developed fever (treated; n = 54) (Fig. 5A). When baseline antibody levels were stratified using median based stratification and assessed by survival analysis, only antibodies to Ripr reached significance (Fig. 5B). Collectively, these results indicate the PCRCR complex is immunogenic and antibodies to PTRAMP-CSS and Ripr may be protective when present in high concentrations.

**Figure 5.**
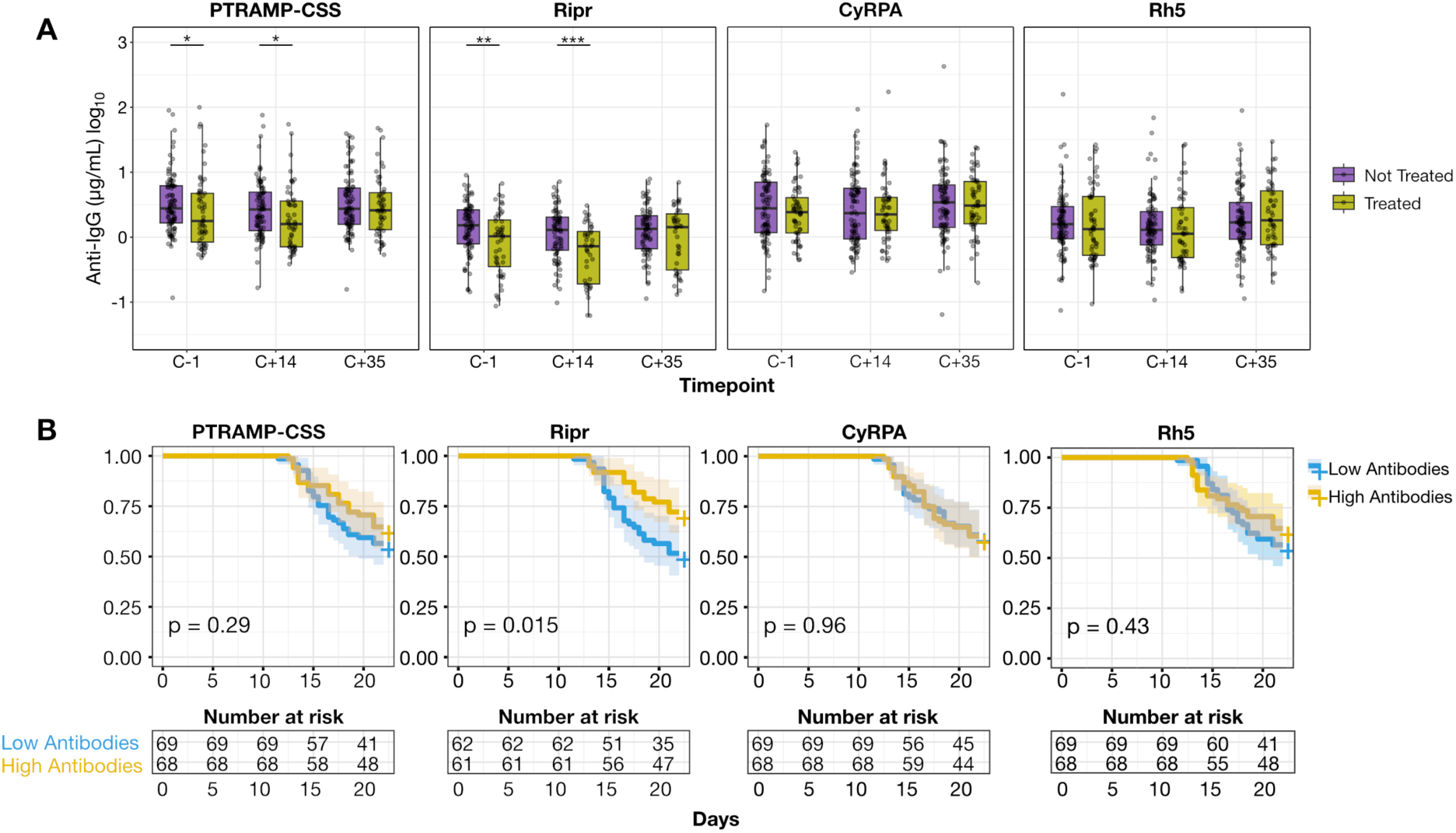
Human IgG antibody responses to PTRAMP-CSS and Ripr may be associated with better clinical outcomes after sporozoite challenge in semi-immune Kenyan adults. **(A)** Antibody kinetics to PTRAMP-CSS, Ripr, CyRPA and Rh5 (left to right) during CHMI-SIKA from 137 individuals. Antibody IgG levels were measured at baseline (one day before challenge (C-1)) and on days 14 and 35 post-challenge (C+14, C+35). Individuals were stratified by whether they received treatment (green, n=54) or did not receive treatment (purple, n=83) which was based on a parasite density or the development of malaria symptoms. Statistical significance between antigen-specific antibody levels and anti-malarial treatment post-challenge was performed using Mann Whitney U test. *p<0.05, ** p<0.01, *** p<0.001. Boxplots show the median and the interquartile ranges. **(B)** Time to treatment survival analysis. Kaplan-Meier curves comparing time to treatment with IgG antibody levels to the PCRCR complex. Survival curves for high antibody responders (blue) and low antibody responders (yellow) are shown. Confidence intervals at 95% intervals are shown as grey shaded bands.

## Discussion

This study provides structural and functional evidence that the PTRAMP-CSS heterodimer, a conserved component of the PCRCR complex (*9*), is a tractable target across multiple *P. falciparum* lifecycle stages. By integrating high-resolution crystal structures, a panel of Nbs and engineered bispecific Fc-fusion proteins with functional assays spanning blood, mosquito and liver stages, we showed that PTRAMP-CSS can be engaged by antibodies to impede merozoite invasion, reduce mosquito infection and limit sporozoite invasion and development in primary human hepatocytes. In addition, higher baseline human IgG titres to PCRCR components were associated with better outcomes after CHMI in a Kenyan cohort, collectively supporting PTRAMP-CSS as a promising antigen for structure-guided multistage vaccine and biologic development.

The crystal structures map both inhibitory and non-inhibitory epitopes across PTRAMP and CSS and reveal a striking spatial determinant of potency with Nbs that recognise epitopes more distal to the merozoite plasma membrane are generally more inhibitory. This pattern was consistent with a model in which the PCRCR complex pre-assembles prior to surface exposure and where steric accessibility and angle of approach determine whether an antibody can effectively block function (*9*, *13*). Mechanistically, the inhibitory Nbs may act by either sterically blocking essential host-parasite interfaces, or constraining conformational mobility required for receptor engagement (*13*, *14*). The discrete, conserved inhibitory epitopes defined on both PTRAMP and CSS therefore provide concrete targets for epitope-focused immunogens and for rational design of multi-specific biologics that simultaneously engage multiple functional surfaces which is an approach that has precedent in structure-guided efforts for Rh5, CyRPA and Ripr (*15*, *19*, *32*, *33*).

A practical advantage of PTRAMP and CSS epitopes is high sequence conservation across global *P. falciparum* isolates. Reformatted bispecific Nb-Fc constructs markedly improved potency versus monovalent Nbs, approaching the *in vitro* activity of leading Rh5 mAbs (*15*). These findings point to two translational paths, either structure-guided immunogen design that biases the humoral response toward conserved inhibitory surfaces to elicit broadly neutralising human antibodies; or antibody-based biologics (bispecific or multispecific formats) as candidate prophylactics or therapeutics. The enhanced potency of multispecific formats supports the latter as a near term biologic approach while vaccine designs are optimised (*28*, *34*).

PCRCR-directed vaccine efforts to date have emphasised Rh5, CyRPA and Ripr, with Rh5 furthest advanced clinically (*20*). Because Rh5 is blood stage restricted, however, it cannot directly provide pre-erythrocytic or transmission blockade (*14*, *18*, *20*). PTRAMP-CSS was expressed in late gametocytes and sporozoites as well as merozoites, offering the possibility of a single-antigen route to multistage coverage. However, the most potent neutralising antibodies reported to date still target Rh5, arguing for combination strategies (for example PTRAMP-CSS with Rh5 or established pre-erythrocytic antigens) to maximise potency and breadth. Structure-guided presentation strategies (epitope scaffolds, VLPs or stabilised complex formats) that mask immunodominant but non-neutralising surfaces such as the Ripr-binding face on CSS should be explored to focus responses onto the inhibitory epitopes identified here (*15–19*, *32*, *33*, *35*).

Although the CHMI-SIKA analyses presented here do not provide unambiguous evidence of association with protection, baseline IgG levels against PTRAMP-CSS and Ripr were higher in the untreated group, and antibodies to Ripr showed associations with protection in the survival analyses. The antibody concentrations reported here are nonetheless considerably lower than those previously linked to immunity. No universal protective threshold has been established for anti-merozoite IgG, as protective concentrations vary by antigen, assay platform, and study population (*36*, *37*) . In children in endemic settings, high-responder status (typically the upper tertile or quartile) is associated with a 30–70% reduction in clinical malaria risk, and breadth of recognition often emerges as a stronger correlate than magnitude (*36*, *38*). For Rh5, the best-characterised PCRCR member, IgG concentrations of approximately 25–100 μg/mL combined with *in vitro* growth inhibitory activity have been associated with *in vivo* protection (*39*, *40*) . Our findings extend this literature by implicating naturally acquired antibodies to PTRAMP-CSS and Ripr in the control of blood stage parasitaemia during CHMI. The borderline significance likely reflects the modest sample size and the use of percentile-based, population-relative cut-offs. Future work should assess whether functional correlates, growth inhibition, opsonic phagocytosis, or complement fixation, which have previously outperformed binding titres (*37*, *41*, *42*), better resolve the link between PCRCR-directed immunity and CHMI outcome.

We define conserved inhibitory epitopes on the PTRAMP-CSS heterodimer, a component of the essential PCRCR invasion complex, and show that nanobodies and engineered bispecific Fc reagents targeting these epitopes potently block *P. falciparum* blood stage invasion, reduce mosquito infection and limit sporozoite invasion of primary human hepatocytes. High-resolution structures reveal sterically accessible, conserved surfaces that can be exploited by structure-guided immunogens or multi-specific biologics, nominating PTRAMP-CSS as a promising single-antigen target for multistage, cross-species malaria interventions.

## Acknowledgements

We thank the Australian Red Cross Lifeblood for the supply of human red blood cells and serum for parasite culture, the WEHI Nanobody Platform for nanobody isolation, and Wai-Hong Tham for the plasmids for nanobody-Fc proteins. We acknowledge Jamie Rossjohn, the Macromolecular Crystallisation Facility, Roxanne Smith, and the Bio21-WEHI Crystallisation Facility where crystallisation screening was undertaken. The authors acknowledge the WEHI Flow Cytometry Facility for their support and assistance. We thank Holger Kanzler, Jacqueline Kirchner, Matthias Paulsner, and Annie Zumsteg from the Gates Foundation for helpful discussion. We thank the surgeons at the Leiden University Medical Center (Stijn van Laarhoven, Sven Mieog and Henk Hartgrink) and our anonymous donors for providing leftover healthy liver resection tissue from which primary human hepatocytes were isolated and tested in the *P. falciparum* sporozoite assays. We acknowledge the flow cytometry and microscopy facilities at the Leiden University Medical Center for access to their equipment. We thank Maureen Blokland-Fink for providing the HepCounter software for the quantification of liver stages. This research was undertaken in part using the MX2 beam at the Australian Synchrotron, part of the Australian Nuclear Science and Technology Organisation (ANSTO) and made use of the Australian Cancer Research Foundation (ACRF) detector. We thank the CHMI-SIKA Study Team in Kenya and the study participants for their collaboration.

## Author Contributions

P.S.L., N.C.J., and R.W.B.C designed experiments, expressed proteins, performed and analysed protein-protein interaction studies. P.S.L. and N.C.J. designed and performed growth inhibition assays and analysed the data. R.W.B.C. raised and identified nanobodies. M.T.N. designed and performed population genetics studies and analysed the data. D.S.M. analysed transcriptomics data.

M.G. and T.R. designed and performed standard membrane feeding assays and analysed the data with P.S.L. M.G. performed and analysed western blots on gametocytes. A.S.P.Y. and J.M.M.K. designed, performed, and analysed data from pre-erythrocytic sporozoite assays. P.S.L. and S.W.S. determined crystal structures and performed structural analyses. D.N. and J.M. performed and analysed human serological assays. A.E.B., F.M.N. M.K, P.B., R.M. A.S.P.Y., A.F.C and S.W.S designed and interpreted experiments.

## Data, code, and materials availability

The crystal structures reported in this manuscript have been deposited in the Protein Data Bank www.rcsb.org (PDB ID codes 25KJ, 25KL, 25KO, 25KQ, 25KR, 25KS). Mini-bulk RNA-seq data used to analyse PCRCR expression profiles in gametocytes have been previously deposited in the NCBI-GEO database under accession GSE314126.

## List of Supplementary Materials

Materials and Methods

Figs. S1 – S8

Tables. S1 – S20

## Materials and Methods

### Ethics

Immunization and handling of the alpaca for scientific purposes was approved by Agriculture Victoria, Wildlife and Small Institutions Animal Ethics Committee, project approval 26-27. Use of human blood and serum for parasite culture was approved by Human Research Ethics Committee (HREC) of Walter and Eliza Institute of Medical Research under approval numbers 19-05VIC-13 and 23-05VIC-09. Primary human liver cells for sporozoite invasion assays were isolated from remnants of surgical material. The samples were anonymously collected and have general approval for their use as granted in accordance with the Dutch ethical legislation (described in the Medical Research Human Subjects Act). It was confirmed by the Medical Ethics Review Committee of Leiden, The Hague and Delft (nWMODIV2_2024030). CHMI-SIKA (ClinicalTrials.gov: NCT02739763) was approved by the KEMRI Scientific and Ethics Review Unit (KEMRI/SERU/CGMR-C/029/3190) and the Oxford Tropical Research Ethics Committee (OxTREC/2-16) and was conducted in accordance with the Declaration of Helsinki (2008).

### Isolation of PTRAMP-CSS specific nanobodies

One alpaca was subcutaneously immunized six times 14 days apart with 100 µg of recombinant PTRAMP-CSS. GERBU FAMA (GERBU Biotechnik GmbH, Heidelberg, Germany) was used as an adjuvant. Blood was collected three days after the last immunization for lymphocyte preparation. Nanobody library construction was carried out according to established methods (*43*). Briefly, alpaca lymphocyte mRNA was extracted and amplified by RT-PCR with a specific primer to generate a cDNA library size of 10_8_ nanobodies with 80% correct sized nanobody insert. The library was cloned into pMES4 phagemid vector amplified in *Escherichia coli* TG1 strain and subsequently infected with M13K07 helper phage for recombinant phage expression.

Phage display was performed as previously described to isolate PTRAMP-CSS Nbs with some modifications (*44*). Phages displaying PTRAMP-CSS Nbs were enriched after two rounds of biopanning against 1 µg of immobilized PTRAMP-CSS protein. After the second round of panning, 94 individual clones were selected for further analyses by ELISA for the presence of PTRAMP-CSS nanobodies. Positive clones were sequenced and annotated using the International ImMunoGeneTics database (IMGT) and aligned in Geneious Prime. A further two rounds of panning were performed against 1 µg of monomeric PTRAMP to identify Nbs specific to PTRAMP. Nb labels begin with a 1, 2 or 3 to distinguish between different biopanning campaigns.

### Nanobody expression and purification

*E. coli* WK6 competent cells were used to express Nbs. Transformed bacteria were grown in Terrific Broth (WEHI Media Kitchen) supplemented with 0.1% (w/v) glucose and 100 µg/mL ampicillin, at 37 °C to an OD_600_ of 0.7, induced with 1 mM IPTG and grown for 16 hrs at 28 °C. The cell pellets were harvested and resuspended in 20% (w/v) sucrose, 20 mM imidazole pH 8.0, 150 mM NaCl DPBS for 15 minutes on ice. 5 mM EDTA was added, incubated for 20 min on ice followed by the addition of 20 mM MgCl_2_ to prevent EDTA chelation, and centrifuged at 20,000 x g for 30 min to pellet the cells and collect the periplasmic extract containing nanobodies. The supernatant was loaded onto 1 – 2 mL Ni-NTA agarose (Qiagen) resin and eluted with PBS pH 7.4, 400 mM imidazole pH 8, 150 mM NaCl, subsequently concentrated and buffer exchanged into PBS.

For single-concentration growth inhibition assays (GIA), Nbs were diluted or concentrated to 5 mg/mL and sterilized using sterile Corning® Costar® 0.22 µm cellulose acetate membrane centrifuge filters. Nbs used for dose dependent GIAs were buffer exchanged into RPMI-1640 medium GlutaMAX HEPES (Gibco) supplemented with 12.5 mg/L hypoxanthine (Sigma), 2 g/L D-glucose, 20 mg/L gentamicin, concentrated to 8 mg/mL, and filter-sterilized as previously described.

### Construct Design

#### CSS

Three glycan deficient versions of CSS (residues 20 to 290, 3D7 clone of *P. falciparum*) were used in this study. The first was subcloned into a modified pTRIEX2 vector with a C-terminal FLAG-tag preceded by a TEV protease cleavage site. This version had four potential N-linked glycosylation sites removed at positions Asn74, Asn192, Asn234 and Asn261 by mutation of Ser76, Ser194, Thr236 and Thr263 to Ala. This construct was used to generate PTRAMP-CSS for immunization studies, BLI and crystallization trials. A second version was subcloned into pAcGP67A vector with a C-terminal FLAG-tag preceded by a TEV protease cleavage site. This construct has all six potential N-linked glycosylation sites removed at positions Asn74, Asn88, Asn192, Asn234, Asn261 and Asn283 by mutation of Ser76, Thr90, Ser194, Thr236 and Thr263 to Ala and Asn283 to Gln. This construct was used to generate PTRAMP-CSS for BLI competition binning experiments as it binds to Ripr with higher affinity (*25*). A monomeric version of CSS was constructed for BLI and crystallization trials. This construct had four glycan sites mutated (Asn74, Asn192, Asn234 and Asn261), Cys30 mutated to an Ala to prevent homodimerization and a C-terminal C-tag (EPEA).

To generate different PfCSS haplotypes, the monomeric CSS construct was modified to include single nucleotide polymorphisms. Certain isolate sequences and strains were chosen to ensure that all possible mutations identified from our population genetics analysis were expressed. The sequences for G224, ML01 and SenT190.08 were sourced from PlasmoDB. Lower frequency mutations were only found in sequences from isolates in the MalariaGen Pf7 database which we have labelled, Ghana-E163V, Guinea-E164K, and Mali-E164G. These isolate sequences were the most common isolates containing amino acid mutations of interests within these countries.

#### PTRAMP

Six glycan deficient versions of PTRAMP were used in this study. The first, comprising residues 42 to 297, was subcloned into the pAcGP67a vector with a C-terminal C-tag. One potential N-linked glycosylation site at Asn195 was removed by mutation of Thr197 to Ala. This construct was used to generate PTRAMP-CSS for immunization studies, BLI and crystallization trials. The second, comprising residues 32 to 309, was subcloned into a modified pTRIEX2 vector with N-terminal SUMO and Flag tags followed by a Tobacco etch virus (TEV) protease cleavage site and a C-terminal Avitag. One potential N-linked glycosylation site at Asn195 was removed by mutation of Thr197 to Ala. This construct was used to measure sera in ELISA. The third, comprising residues 31 to 307, was subcloned into the pAcGP67a vector with a C-terminal C-tag. Four potential N-linked glycosylation sites were removed, at positions Asn112, Asn149 and Asn155 by mutation to Gln, and at position Asn195 by mutation of Thr197 to Ala. This construct was used for BLI competition studies, as it binds to Ripr with higher affinity (*25*). A monomeric version of PTRAMP was constructed for Nb panning, BLI and crystallization trials. Comprising residues 42 to 297, this construct had one potential glycan site at Asn195 removed by mutation of Thr197 to Ala, Cys60 mutated to a Ser and a C-terminal C-tag (EPEA). Finally, the PTRAMP growth factor (residues 75 to 241) and TSR (residues 242 to 297) domains were subcloned into the pAcGP67A vector with a C-terminal C-tag. Again, one potential N-linked glycosylation site at Asn195 was removed by mutation of Thr197 to Ala in the GFD construct. These constructs were used to generate PTRAMP subdomains for BLI and crystallography studies.

#### Ripr

The gene for Ripr (residues 20 to 1086) was subcloned into the pAcGP67A vector with a C-terminal His-tag (*9*). Additionally, a construct with an N-terminal Avitag was generated for ELISAs and BLI competition binning experiments. To generate Ripr-tail, residues 717 to 1086 of Ripr were subcloned into the pAcGP67A vector with a C-terminal His-tag. Two potential N-linked glycosylation sites were removed by mutation of Asn964 and Asn1021 to Gln. This construct was used for BLI competition studies.

#### CyRPA

The gene for CyRPA (residues 29 to 362) was subcloned into a modified pcDNA3.4-TOPO plasmid with an N-terminal IL-2 signal sequence and a C-terminal Flag tag preceded by a TEV protease cleavage site. Three potential N-linked glycosylation sites at Asn145, Asn322 and Asn338 were removed by mutation of the glycan site Thr or Ser residues to Ala. To generate a biotinylated CyRPA, the TEV protease cleavage site was replaced with an Avitag.

#### Rh5

The gene for PMX cleaved PfRh5 (residues 145 to 526) was subcloned into pAcGP67A with a C-terminal C-tag. Three potential N-linked glycosylation sites as Asn214, Asn284 and Asn297 were removed by mutation of the glycan site Thr or Ser residues to Ala. To generate a biotinylated Rh5, the TEV protease cleavage site was replaced with an Avitag.

#### Nb-Fc and BsNb-Fc constructs

Nb-Fc constructs were generated as previously described (*34*). Briefly, nanobody-encoding sequences were amplified by PCR and inserted into pHLSec-Fc via *Age*I and *Nhe*I so that the nanobody was genetically fused to human IgG1-Fc backbone. Similarly, BsNb-Fc genes were synthesized by Twist Biosciences and genetically fused to IgG1-Fc backbone.

#### Recombinant protein expression and purification

PTRAMP, CSS, PTRAMP-CSS, Ripr and Rh5 constructs were expressed in Sf21 cells and secreted into the medium as soluble protein. The supernatant was harvested after 3 days of incubation and purified by either CaptureSelect™ C-tag XL Affinity resin (ThermoScientific), M2 Anti-Flag Affinity Resin (Sigma), or Ni-NTA agarose (Qiagen) resin, followed by size exclusion chromatography (S200 Increase 10/300 GL, Cytiva). To remove N-linked glycans prior to crystallization, PTRAMP_75-241_ was pre-treated with PNGaseF-GST in a 1:1 molar ratio and incubated overnight at 37 °C. Excess PNGaseF-GST was then removed by running the reaction over GST agarose (Sigma) prior to co-complexation with Nbs.

CyRPA was expressed via transient transfection of Expi293F cells, and soluble protein was purified from the culture medium via M2 Anti-Flag Affinity Resin (Sigma) followed by size exclusion chromatography (S200 Increase 10/300 GL, Cytiva).

Nb-Fcs and BsNb-Fcs were expressed via transient transfection of Expi293F cells, and soluble protein was purified from the culture medium via HiTrap MabSelect Prism A^TM^ column, followed by size exclusion chromatography (S200 Increase 10/300 GL, Cytiva or HiLoad 16/600 Superdex 200, Cytiva) with PBS.

Proteins were biotinylated according to established protocols (*45*).

#### Biolayer interferometry studies

To determine epitope bins and binding kinetics of Nbs to PTRAMP-CSS, biolayer interferometry experiments were conducted at 25 °C in solid black 96-well plates agitated at 1000 rpm with kinetics buffer (PBS pH 7.4, 0.1% (w/v) BSA, 0.02% (v/v) Tween-20) using an Octet RED96e instrument (Sartorius).

For binding kinetic studies, His-tagged Nbs were diluted in kinetics buffer at 10 µg/mL and immobilised onto Ni-NTA sensors (Sartorius). Following a 60 sec baseline step, biosensors were dipped into wells containing a two–fold dilution series of PTRAMP-CSS from 500 – 15.63 nM. Sensors were then dipped into wells containing buffer only to measure the dissociation rate. Nbs were also tested against monomeric PTRAMP and CSS, and binding to either the TSR or Growth Factor Domain of PTRAMP was measured for PTRAMP-specific nanobodies using similar methods. Kinetics curves were fitted to a 1:1 model, the K_D_ reported is the mean of two independent experiments.

Similar to binding studies, for epitope binning experiments, 5 – 10 µg/mL of primary Nbs were immobilized onto Ni-NTA sensors and after a baseline step in buffer, remaining binding sites on Ni-NTA biosensors were quenched with H2, an anti-PTRAMP Nb that cannot bind to the PTRAMP-CSS heterodimer. The sensors were then dipped into wells containing 250 nM of PTRAMP-CSS until the Nbs were fully saturated. Following another baseline step, the biosensors were then dipped into secondary Nb to determine competition binding. Data presented in Supplementary Figure 2 represent the percent competing nanobody binding compared with the maximum competing response. A cut-off of less than 33 was considered competing.

To determine if PTRAMP or CSS Nbs were able to compete with Ripr binding onto PTRAMP-CSS, 20 µg/mL of biotinylated Ripr was loaded onto SAX2 sensors, dipped into PTRAMP-CSS (with four and six glycans removed, respectively, to improve affinity to Ripr) until saturated, and after a baseline step, dipped into Nbs to determine binding.

### Parasite, insect and mammalian cell culture

#### Culture of asexual stage *P. falciparum* parasites

3D7 *P. falciparum* parasites were obtained from David Walker, Edinburgh University and were routinely cultured as previously described (*46*). Asexual stage parasites were cultured in human O^+^ erythrocytes (Australian Red Cross Bloodbank, South Melbourne, Australia) at 4% haematocrit (HCT) in complete Roswell Park Memorial Institute (RPMI) media which consisted of RPMI-1640 medium GlutaMAX HEPES (Gibco) supplemented with 12.5 mg/L hypoxanthine (Sigma), 2 g/L D-glucose, 20 mg/L gentamicin, 0.25% (w/v) Albumax II (Gibco) and 5% (v/v) heat-inactivated human serum (Australian Red Cross Bloodbank, South Melbourne, Australia). Cultures were incubated at 37 °C in a gaseous mix of 1% O_2,_ 5% CO_2_ in 94% N_2._ All cultures were sub-cultured every two days with new culture media and erythrocytes to maintain 4% HCT. Parasites were synchronized at ring-stages with 5 x culture volume of 5% w/v sorbitol. Trophozoite stage parasites (>16 hrs post-invasion) were harvested for growth inhibition assays.

#### *P. falciparum* gametocyte culture

Asexual stage of NF54/iGP2 (*47*) parasites were maintained routinely as outlined above, except that the culture media was supplemented with 2.5 mM D-(+)-Glucosamine hydrochloride (GlcN). To induce gametocyte conversion, 30 mL culture of synchronised ring stage parasites (1-3%) were washed off GlcN with gametocyte culture media (RPMI-1640 medium GlutaMAX HEPES supplemented with 12.5 mg/L hypoxanthine, 2g/L D-glucose, and 10% (v/v) heat-inactivated human serum) and allowed to reinvade for 48 hrs. Committed ring stages (Day 0) were then transferred into a 6-well plate (Corning) and fed daily until Day 12.

### Growth inhibition assays

One-cycle growth inhibition assays were mostly performed as previously described (*18*). Briefly, 5 µL of anti-PTRAMP-CSS Nbs or 1G12 as a positive control anti-Ripr mAb were added to 96-well round bottom plates followed by 45 µL of trophozoite-stage parasites at 0.4 % parasitemia and 2% HCT, for a final nanobody concentration of 0.5 mg/mL. For titration experiments, Nbs were first buffer exchanged into incomplete RPMI medium (supplemented only with 0.2% NaHCO_3_, 0.37 mM hypoxanthine and 31.25 µg/mL gentamicin) and 50 µL of Nb was used for a two-fold twenty-step dilution series. 25 µL of 0.8% parasitemia parasites at 4% HCT (supplemented with double the concentration of Albumax and HIHS Serum) were added to wells. After incubation for 48 hr, each well was fixed at RT for 30 min with 50 µL of 0.25% v/v glutaraldehyde (ProSciTech) diluted in PBS. Following centrifugation, the supernatant was discarded, the cells washed in PBS and stained with 50 µl of SYBR Green (Invitrogen) diluted in PBS. The parasitaemia of each well was determined by counting 100,000 cells by flow cytometry using an Attune NxT Flow Cytometer (ThermoFisher) or Novocyte Flow Cytometer (Agilent). Growth was expressed as a percentage of parasitaemia obtained using a non-immune IgG or vehicle control. All samples were tested in triplicate and standard error of the mean calculated.

Synergy GIAs of all pairs of inhibitory nanobodies were assessed as described in (*27*). The growth inhibition activity of one nanobody was held constant at approximately 20% GIA (1); and a second nanobody (2) across a two-fold 20-step dilution curve beginning at 1 mg/mL; and (3) the combination of the first nanobody held at a constant concentration with the second nanobody across its dilution curve. The bliss additivity was determined based on the measured activity from each antibody alone (1 and 2) using the following formula from Ragotte et al., (*27*):

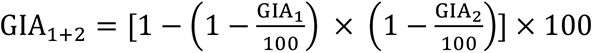

### Three-dimensional structure determination of PTRAMP-CSS-Nb, PTRAMP-Nb and CSS-Nb complexes

To co-complex Nbs with either PTRAMP-CSS, CSS, PTRAMP, TSR domain of PTRAMP or GFD of PTRAMP, the antigens and Nbs were mixed in a 1:3 molar ratio in 20 mM HEPES pH 7.5 150 mM NaCl, and excess Nb was purified away via size exclusion chromatography (Superdex 200 Increase 10/300 GL, Cytiva or Superdex 75 Increase 10/300 GL, Cytiva, depending on size of equimolar Nb-antigen complex). The complexes were concentrated to 5.0 mg/mL, mixed 1:1 with mother liquor, and set up in large-scale, sparse-matrix sitting-drop crystallization trials performed at either the WEHI/Bio21 Crystallization Facility (Parkville, Australia) or the Monash Macromolecular Crystallization Facility (Clayton). Crystals of 1H10-CSS and 2C2-PTRAMP were harvested directly from the sitting-drop plates, whereas 1G8-PTRAMP-CSS, 1G2-CSS, 3E2-PTRAMP-GFD and 2D11-3F4-PTRAMP-TSR were further refined in-house by hanging drop vapor diffusion crystallization trials.

2C2-PTRAMP crystallized in 0.2 M sodium thiocyanate (KSCN), 20% (v/v) PEG 3350 and were cryoprotected in mother liquor containing 15% (v/v) ethylene glycol. 3E2-PTRAMP-GFD crystallized in 20% (v/v) PEG 8K, 0.1 M CHES pH 9.5 after seeding with crystals grown in 0.1 M CHES pH 9.0, 20% (v/v) PEG 8K, after 36 hours, and were cryoprotected in 20% (v/v) glycerol. Crystals of 2D11-3F4-PTRAMP-TSR were obtained in 0.1 M bis-tris-propane (BTP) pH 5.5, 20% (v/v) PEG 3350, after 24 hrs, and were cryoprotected in 20% (v/v) glycerol. 1G2-CSS crystallized in 0.1 M BTP pH 6.5, 0.2 M KSCN and 16% (v/v) PEG 3350, after 24 hrs, and were cryoprotected in 20% (v/v) ethylene glycol. 1H10-CSS crystallized in 0.1 M HEPES pH 7.0, 0.2 M calcium chloride and 20% (v/v) PEG 8K, after 24 hrs, and were cryoprotected in 20% (v/v) glycerol. Crystals of 1G8-PTRAMP-CSS were grown in 0.1 M BTP pH 7.5, 0.2 M sodium/potassium phosphate and 15% (v/v) PEG 3350, and cryoprotected in 20% (v/v) ethylene glycol.

Diffraction data were collected at the MX2 beamline at the Australian Synchrotron (Clayton, Victoria), processed and merged using the XDS (*48*) and Aimless (*49*) packages. Diffraction data for the 1H10-CSS complex were anisotropic and were therefore processed using Staraniso (*50*). CSS from the H2-PfCSS crystal structure (PDB: 7UNZ, (*9*)) was first used as a model for molecular replacement of CSS in the crystal structures of 1G2-CSS, 1H10-CSS and 1G8-PTRAMP-CSS. The position of 1G2, 1H10 and 1G8 Nbs in the 1G2-CSS, 1H10-CSS and 1G8-PTRAMP-CSS crystal structures were determined by molecular replacement using the structure of nanobodies from Nanobody 25 from 5O8F (*51*), Nanobody 2 from 5JA8 (*52*), and Nanobody 22 from 5LHR (*53*), respectively, with their CDR3s removed. The models used to position 2C2, 3E2, 2D11 and 3F4 on PTRAMP were Nanobody 22 from 5LHR (*53*), Nanobody F3 from 8E0E (*54*) and Nanobody 25 from 5O8F (*51*) and Nanobody 14 from 5JMR (*55*) respectively. An AlphaFold2 model of PTRAMP (Q8I5M8-F1) was used for molecular replacement of 2C2-PTRAMP which had a resolution of 1.5 Å. This high-resolution structure was subsequently used as model for molecular replacement of the TSR domain (242 – 297) in the 2D11-3F4-PTRAMP-TSR crystal structure and the GFD domain (75 – 241) in the 3E2-PTRAMP-GFD structure. All molecular replacements were determined by Phaser (*56*). Structures were manually improved in Coot (*57*) and refined in Phenix.refine (*58*) in an iterative process. Interactions between nanobodies and antigens were determined by PISA and figures were prepared using ChimeraX (*59*). The Dali server was used to analyse the structural homology of PTRAMP (*60*).

### Structure prediction

The AlphaFold3 server was used to predict the structure of the PTRAMP, CSS and Ripr complex (*23*).

### Genomic data and quality control

We analysed *P. falciparum* whole-genome sequencing data from 20,646 field isolates in the MalariaGEN Pf7 open-access dataset (*26*) . Reads were mapped to the *P. falciparum* 3D7 v3 reference genome and variants were jointly genotyped using GATK v4 as previously described (*61*) . Analyses were restricted to the core genome, excluding subtelomeric, internal hypervariable, centromeric, mitochondrial, and apicoplast regions. We retained only SNPs passing all quality filters, with read depth ≥5 and mapping quality QUAL ≥50 and excluded loci with >10% missing genotypes in the nuclear genome. Chromosome-level Variant Call Format (VCFs) were merged using bcftools after removing indels and complex alleles (*62*) . Samples with >20% missing genotypes across filtered SNPs were excluded, and only PASS-annotated variants were retained. Multiplicity of infection (MOI) was estimated using DEploid and moimix (available at https://bahlolab.github.io/moimix/), with population allele frequencies calculated at the WHO-defined regional level (*63*) . Samples with MOI = 1 or dominant clone proportion >95% were treated as clonal. For MOI ≥2 samples, minimum read depth thresholds of 20× (MOI = 2) and 30× (MOI >2) were required for haplotype phasing. Clonal and dominant-clone (>65%) MOI-2 samples were phased using allelic depth; balanced MOI-2 samples (50:50) were phased using SHAPEIT4 on the full high-quality callset. Major-clone haplotypes for MOI ≥3 samples were derived from high-depth sites. After phasing, variants with >10% missingness were further removed, consistent SNP identifiers were assigned.

Target antigen sequences were extracted from the final call set using dominant clones and converted from VCF to FASTA format. Singletons and putative sequencing artefacts were removed prior to analysis. Diversity metrics i.e. nucleotide or sequence diversity (π), number of segregating sites, and synonymous and non-synonymous SNP counts were calculated using the *VaxPack* R package (https://github.com/BarryLab01/vaxpack). Sequence diversity values were mapped onto three-dimensional protein structures using the biostructmap Python module (*64*). Sequence logos were generated using the WebLogo command-line tool (*65*).

### Transcriptomic analysis of PCRCR expression during gametocyte development

Transcriptomic data of synchronous day 3,6,9 and 12 NF54/iGP2 parental strain parasites was analysed for the expression of PCRCR components alongside the stage specific control as previously described (*66*).

### Standard Membrane Feeding Assays

Female *Anopheles stephensi* mosquitos were fed with 0.5 mL blood meals consisting of 50/50 parasitised RBCs at 0.5% stage V gametocytemia and heat-inactivated O^+^/A^+^ human serum at 37 °C. Blood meals were prepared as a batch and split equally into separate tubes, where each blood meal was combined with either 10 µL Nanobody-Fcs (5 mg/mL), 10 µL TB31F (5 mg/mL), or 10 µL 1X DPBS. Mosquitoes were allowed to feed for 1-2 hours in the dark. Mosquito midguts (20-30 from each group) were dissected 8 days post-feeding and stained with 0.1% mercurochrome to visualise oocysts. Oocyst numbers were counted manually, then plotted onto a graph and analysed with GraphPad Prism 10.1.1 (GraphPad Software, inc).

### Western blots

Schizont and gametocyte (both NF54/iGP2 line) lysates were extracted from saponin lysed pellets using 1X TNET buffer (1% Triton X-100, 150 mM NaCl, 10 mM EDTA, 50 mM Tris pH 7.4) prior to mixing with non-reducing sample buffer. 15 µL lysates were loaded onto 4-12% Bis-Tris Gel (Invitrogen), electrophoresed at 180V for 45 minutes in 1X MES Buffer, and blotted onto a nitrocellulose membrane using iBlot 2 system (Invitrogen) according to the manufacturer’s instructions. The membrane was blocked with 5% (w/v) skim milk diluted in 1X PBS-T overnight at 4 ^°^C.

Salivary gland sporozoites (NF54/iGP2 line) were obtained by dissecting infected *A. stephensi* mosquitoes 18 days post-feeding. Sporozoite suspension (in 1X DPBS) was then lysed by boiling in non-reducing sample buffer. Lysates equivalent to 200,000 sporozoites (with the addition of 100 mM DTT) were electrophoresed, blotted, and blocked as described above.

Primary antibody/nanobody was used at a following concentration: 20 µg/mL HRP-conjugated Nanobody-Fcs; 1/20000 rabbit α-HSP70; 1/2000 mouse α-Pfs16 (mAb B4); 1/1000 rat α-CSS (mAb 2D2); and 1/1000 mouse α-PTRAMP (mAb 1D9). Secondary goat α-rabbit IgG (HRP), goat α-mouse IgG (HRP), and goat α-rat IgG (HRP) were used at 1/2000.

### Sporozoite Traversal Assay

HC-04.J7 cells were maintained in DMEM:F12 1:1 (Gibco) supplemented with 10% Hi-FCS and 100 units/mL Penicillin-Streptomycin prior to infection. HC-04.J7 cells were seeded in a collagen-coated 96-well plate at a 40,000 cells/well density one day prior to infection.

Sporozoite traversal assays were performed as previously described (*67*). Briefly, sporozoites were isolated from NF54-infected *A. stephensi* mosquitoes (day 17-19 post-bloodmeal) into ice-cold William’s B media (*68*). Sporozoite lysates were pre-mixed with individual nanobodies (100 µg/mL or 500 µg/mL) together with FITC-conjugated dextran (1 mg/mL, 10,000 MW, Sigma Aldrich: FD10S-100MG) and 10% human serum. Upon sporozoite addition (50,000/well), cells were incubated for 4 h at 37 °C with 5% CO_2_. After incubation, cell suspensions were analysed using flow cytometry. The percentage of dextran-FITC^+^ HC-04.J7 cells was detected on a BD FACSCanto II. Data were analysed using FlowJo v11.

### Sporozoite Developmental Assay

Primary human hepatocytes isolated from liver-resections (*68*) were plated in a collagen-coated 96-well plate at 62,500 cells/well density. Cells were maintained in William’s E media supplemented 1x B27, 100 units/ml of Penicillin-Streptomycin and 0.5 µM of IWP2 (*69*). Infectious NF54 sporozoites were isolated as described above and were subsequently pre-mixed with individual nanobodies (100 µg/mL) and 10% heat-inactivated human sera. Sporozoites were added in triplicate (50,000/well). Infected cells were maintained for 3 days with daily media changes, until fixing with 4% PFA for 10 minutes. Cells were permeabilised with 1% Triton-X100 and blocked with 3% BSA. Parasites were detected using anti-GAPDH antibodies (1:25000; European Malaria Reagent Repository Catalogue number 7.2). Secondary goat anti-mouse Alexa Fluorophore 488 (Thermofisher: A11029) was used at 1:300 dilution. Nuclei were stained with DAPI (Thermofisher: D1306) at 300 nM. Individual wells were imaged using the ImageXpress high-throughput confocal microscope (Molecular Devices) at 20x magnification. Infected cells were quantified using in-house HepCounter software (v4.9).

### Controlled Human Malaria Infection in Semi-immune Kenyan Adults (CHMI-SIKA)

We analysed samples from the CHMI-SIKA study (*70*), an open-label, non-randomised study of healthy adults recruited from areas of moderate-to-high (Ahero), moderate (Kilifi South), and low (Kilifi North) *P. falciparum* transmission. All participants were *P. falciparum*-negative at baseline. A total of 161 participants were intravenously challenged with 3,200 cryopreserved NF54 *P. falciparum* sporozoites (Sanaria PfSPZ Challenge); 19 were excluded from analysis owing to significant baseline antimalarial drug levels, leaving 142 in the primary analysis. Samples from 137 of these participants were available for the present study.

Following challenge, participants were monitored for 35 days, during which parasite growth and clinical symptoms were assessed and immunological samples collected. *P. falciparum* parasitaemia was monitored by quantitative PCR (qPCR) twice daily from days 7–14 and once daily from days 15–21. Participants were classified as “treated” or “untreated” according to a predefined clinical protocol, with treatment indicated by either a parasite density exceeding 500 parasites/µL or the development of malaria symptoms. All participants received artemether–lumefantrine on meeting treatment criteria or at study completion (day 21). This classification served as a proxy for the participant’s ability to control parasite growth and clinical disease.

### ELISA

High binding ELISA plates (NUNC, Maxisorp, Thermo Scientific) were coated with recombinant Rh5, Ripr, CyRPA and PTRAMP-CSS, at a final concentration of 125 nM in PBS for 1 hour at room temperature. Plates were washed three times in PBS-T and blocked with 4% skim milk in PBS-T for one hour, followed by an additional three washes in PBS-T. Heat-inactivated serum samples were diluted 1:100 in PBS and added to the wells. Plates were incubated for one hour at RT and washed. HRP-conjugated anti-human secondary antibody diluted 1:5000 in PBS-T was added and incubated for one hour. Plates were washed and developed with 50 µL of o-phenylenediamine (OPD) substrate for 20 minutes at RT in the dark. The reaction was stopped with 2 M of sulfuric acid per well. Optical density was measured at 492 nm using the Synergy™ plate reader (Biotek Instruments Inc.). All assays were performed in duplicate, and assay was repeated if coefficient of variation between duplicates exceeded 20%. Serial dilutions of monoclonal antibodies were included on each plate as standards to quantify antibody concentration. Pooled hyperimmune sera from individuals residing in a malaria endemic area were included as a positive control.

## Statistical Analysis

Statistical analysis was done using Graphpad Prism v 9.1.0 for Mac. EC_50_ for GIA curves was determined through a four-parameter (max, min, variable Hill slope, EC_50_) logistic regression with the upper bound constrained to 100% GIA. A *p* value < 0.05 was considered significant. Human antibody data was analyzed using R v 4.6.0.

