## Supplementary Materials for "Targeting PTRAMP-CSS potently inhibits *P. falciparum* across blood, liver and mosquito stages"

Pailene S. Lim *et al.*

Corresponding authors:

Stephen W. Scally,

Alan F. Cowman,

**This PDF includes:**

Figs. S1 to S8

Tables S1 to S20

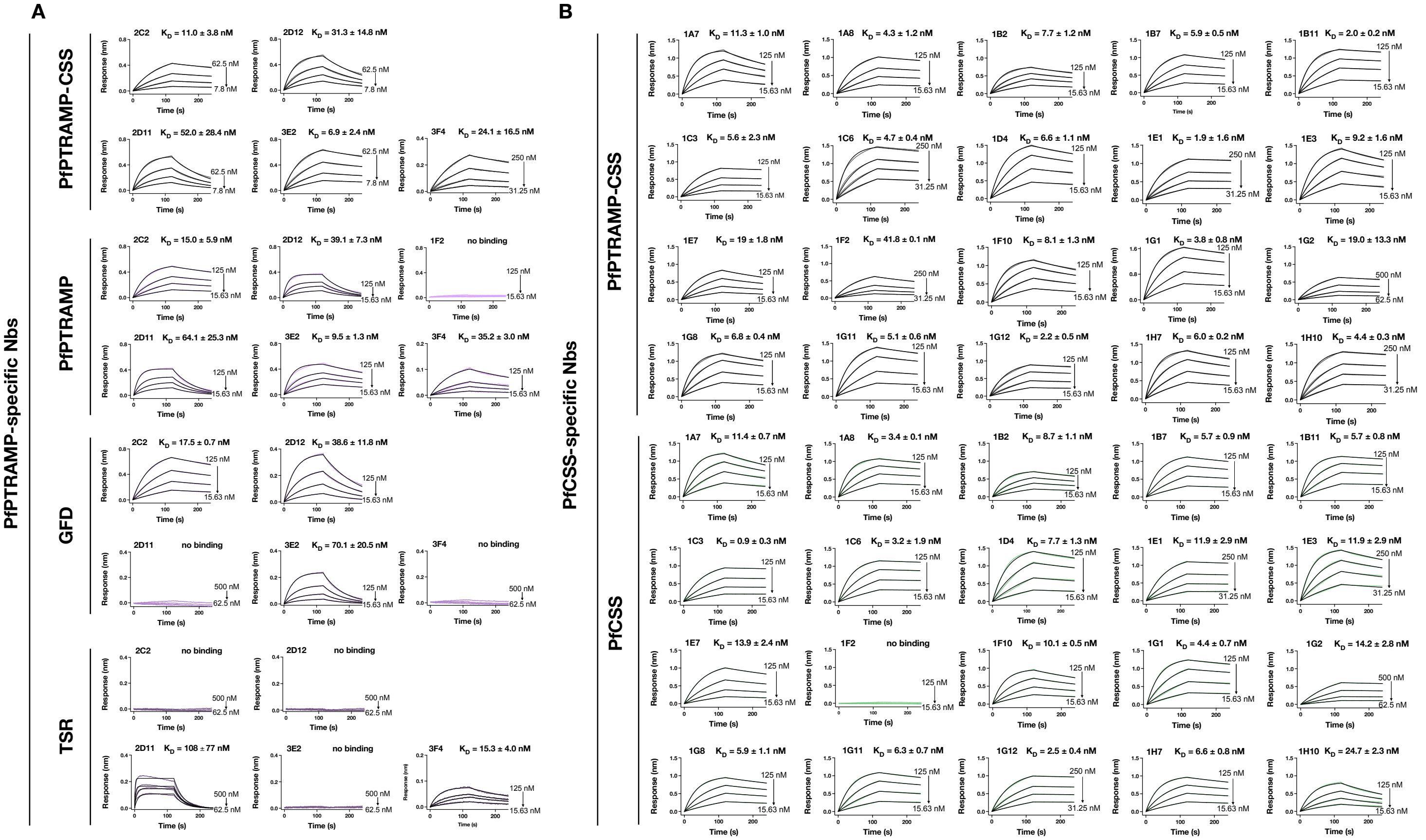


**Supplementary Figure 1. Binding kinetics of nanobodies to PfPTRAMP, CSS or PfPTRAMP-CSS.** **(A)** Representative biolayer interferometry sensorgrams of α-PfPTRAMP nanobodies to PfPTRAMP-CSS, PfPTRAMP and the growth factor domain (GFD) and thrombospondin type 1 repeat (TSR) domain. Dilution series data are shown in color for each protein and 1:1 model best fit is shown in black. Starting and end concentration of two-fold dilution series are also shown for each nanobody. **(B)** Representative sensorgrams of α-PfCSS nanobodies binding to PfPTRAMP-CSS and PfCSS. Dilution series data is shown in grey for PTRAMP-CSS and green for PfCSS. See also table S1 and S2. $K_D$ (nM) ± standard error of the mean is representative of two independent biological binding assay experiments.

**A**

Nanobody 1 (on sensor) × Nanobody 2 (in well)

| Nanobody 1 (on sensor) | 1A7 | 1F10 | 1H7 | 1B7 | 1G1 | 1B2 | 1B11 | B6 | 1G11 | 1G12 | 1D4 | 1E3 | 1C3 | 1E7 | 1C6 | 1A8 | B1 | 1G8 | D2 | 1G2 | 1H10 | 1E1 | 1F2 |
|---|---|---|---|---|---|---|---|---|---|---|---|---|---|---|---|---|---|---|---|---|---|---|---|
| 1A7 | 0 | 0 | 0 | 0 | 0 | 0 | 0 | 0 | 0 | 0 | 0 | 0 | 34 | 32 | 27 | 25 | 58 | 50 | 35 | 41 | 56 | 41 | 57 |
| 1F10 | 0 | 0 | 0 | 0 | 0 | 0 | 0 | 0 | 0 | 0 | 0 | 0 | 52 | 61 | 3 | 57 | 43 | 0 | 18 | 44 | 38 | 43 | 50 |
| 1H7 | 0 | 0 | 0 | 0 | 0 | 0 | 0 | 0 | 0 | 0 | 0 | 0 | 0 | 5 | 0 | 0 | 4 | 55 | 36 | 54 | 54 | 59 | 62 |
| 1B7 | 26 | 9 | 31 | 1 | 0 | 0 | 0 | 0 | 0 | 0 | 0 | 0 | 69 | 65 | 75 | 100 | 72 | 0 | 68 | 77 | 74 | 67 | 87 |
| 1G1 | 0 | 12 | 9 | 0 | 0 | 0 | 0 | 0 | 0 | 0 | 0 | 0 | 76 | 100 | 85 | 81 | 65 | 0 | 73 | 96 | 86 | 88 | 94 |
| 1B2 | 5 | 19 | 36 | 16 | 0 | 0 | 0 | 0 | 0 | 0 | 0 | 0 | 51 | 59 | 0 | 85 | 86 | 93 | 95 | 84 | 93 | 100 | 92 |
| 1B11 | 5 | 39 | 44 | 30 | 17 | 0 | 13 | 15 | 21 | 11 | 0 | 0 | 100 | 86 | 1 | 74 | 77 | 70 | 74 | 82 | 85 | 82 | 87 |
| B6 | 0 | 0 | 0 | 0 | 0 | 0 | 0 | 0 | 0 | 0 | 0 | 0 | 0 | 0 | 0 | 0 | 19 | 69 | 63 | 65 | 72 | 82 | 79 |
| 1G11 | 0 | 3 | 15 | 0 | 0 | 0 | 0 | 0 | 0 | 0 | 0 | 0 | 0 | 0 | 0 | 0 | 0 | 75 | 59 | 63 | 75 | 77 | 92 |
| 1G12 | 6 | 23 | 21 | 10 | 11 | 0 | 1 | 0 | 7 | 27 | 0 | 0 | 0 | 0 | 0 | 0 | 0 | 64 | 49 | 71 | 66 | 72 | 78 |
| 1D4 | 0 | 17 | 33 | 3 | 0 | 0 | 0 | 0 | 0 | 0 | 0 | 0 | 0 | 0 | 0 | 0 | 23 | 96 | 90 | 99 | 100 | 94 | 96 |
| 1E3 | 0 | 0 | 5 | 0 | 0 | 0 | 0 | 0 | 0 | 0 | 0 | 0 | 0 | 0 | 0 | 0 | 0 | 100 | 82 | 94 | 88 | 99 | 100 |
| Ripr | 36 | 33 | 21 | 23 | 0 | 0 | 30 | - | 5 | 9 | 20 | 2 | 43 | 40 | 30 | 13 | - | 11 | 86 | 74 | 71 | 63 | 76 |
| 1C3 | 71 | 80 | 79 | 89 | 79 | 71 | 99 | 30 | 0 | 0 | 30 | 0 | 12 | 0 | 25 | 9 | 45 | 72 | 86 | 89 | 72 | 84 | 74 |
| 1E7 | 80 | 97 | 90 | 87 | 96 | 76 | 88 | 0 | 0 | 0 | 0 | 0 | 0 | 0 | 0 | 0 | 38 | 89 | 100 | 100 | 73 | 98 | 86 |
| 1C6 | 65 | 63 | 50 | 75 | 71 | 0 | 27 | 0 | 0 | 0 | 0 | 0 | 0 | 0 | 6 | 0 | 30 | 73 | 63 | 73 | 66 | 71 | 70 |
| 1A8 | 76 | 93 | 72 | 75 | 88 | 66 | 74 | 0 | 0 | 0 | 0 | 0 | 0 | 0 | 6 | 0 | 31 | 74 | 80 | 90 | 71 | 97 | 76 |
| B1 | 40 | 53 | 45 | 59 | 51 | 38 | 55 | 35 | 0 | 0 | 0 | 0 | 0 | 0 | 8 | 0 | 18 | 47 | 65 | 68 | 53 | 75 | 60 |
| 1G8 | 59 | 59 | 79 | 0 | 21 | 58 | 64 | 80 | 78 | 75 | 68 | 73 | 60 | 69 | 79 | 64 | 57 | 0 | 49 | 54 | 60 | 77 | 69 |
| D2 | 43 | 94 | 81 | 76 | 84 | 41 | 67 | 98 | 72 | 60 | 55 | 59 | 63 | 78 | 87 | 49 | 100 | 83 | 52 | 24 | 18 | 39 | 49 |
| 1G2 | 78 | 94 | 89 | 87 | 88 | 69 | 72 | 88 | 79 | 77 | 65 | 66 | 62 | 92 | 85 | 75 | 86 | 76 | 66 | 30 | 22 | 48 | 60 |
| 1H10 | 100 | 100 | 100 | 100 | 100 | 100 | 100 | 100 | 100 | 100 | 100 | 100 | 79 | 96 | 100 | 90 | 77 | 95 | 82 | 0 | 0 | 27 | 46 |
| 1E1 | 67 | 96 | 87 | 92 | 91 | 73 | 88 | 77 | 73 | 71 | 81 | 79 | 78 | 93 | 98 | 61 | 66 | 92 | 62 | 15 | 6 | 27 | 43 |
| 1F2 | 35 | 52 | 56 | 48 | 40 | 34 | 39 | 39 | 61 | 51 | 34 | 30 | 34 | 44 | 48 | 36 | 40 | 35 | 37 | 0 | 0 | 4 | 24 |

**B**

Nanobody 1 (on sensor) × Nanobody 2 (in well)

| Nanobody 1 (on sensor) | 2C2 | 2D12 | 3E2 | 3F4 | 2D11 |
|---|---|---|---|---|---|
| 2C2 | 15 | 23 | 94 | 100 | 78 |
| 2D12 | 14 | 23 | 91 | 99 | 76 |
| 3E2 | 62 | 68 | 32 | 85 | 67 |
| 3F4 | 42 | 48 | 61 | 31 | 48 |
| 2D11 | 67 | 76 | 87 | 89 | 22 |
| Ripr | 71 | 64 | 100 | 77 | 75 |

**Supplementary Figure 2. Epitope binning of CSS and PTRAMP nanobodies. (A)** Competition binning of α-PfCSS nanobodies against PfCSS and biotinylated PfRipr. **(B)** Competition binning of α-PfPTRAMP nanobodies against PfPTRAMP-CSS and PfRipr. Primary nanobodies or antigen bound to the biolayer interferometry sensor are shown on the left-hand side whilst secondary nanobodies or antigen are listed on the top of the tables. Data indicates the percent of competing nanobody or PfRipr compared to the maximum competing nanobody response for each column (secondary nanobody). A cut-off of >33% competition was used to determine if nanobodies were competing (red) or non-competing (blue). Nanobodies are colored according to their epitope bins shown in Figure 1A. B1, B6 and D2 were previously identified epitopes from Scally *et al.* (1).

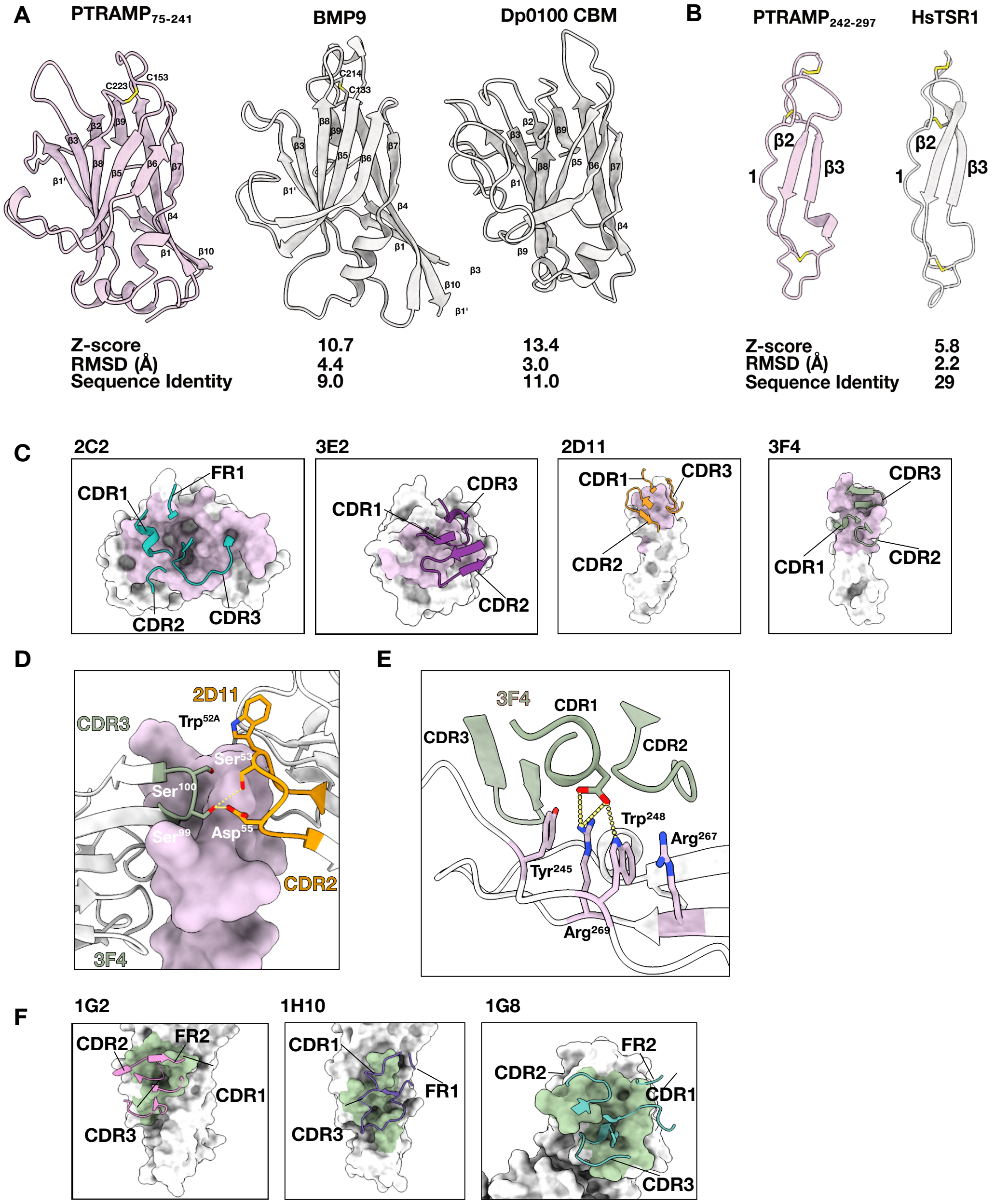


**Supplementary Figure 3. Structural and biophysical analysis of nanobody binding to PTRAMP and CSS. (A)** Comparison of PTRAMP GFD with previously solved crystal structures of BMP9, part of the TGF-β superfamily and carbohydrate binding module (CBM) of Dp0100, a carbohydrate binding protein. **(B)** Comparison of PTRAMP TSR domain with previously solved crystal structure of human TSR1 domain (HsTSR1). Disulfide bonds are shown in yellow. Z-score, RMSD (Å), and sequence identity are shown below. **(C)** Interactions between PTRAMP and α-PTRAMP Nbs. Epitopes are colored in pink and interacting CDRs or FRs of Nbs are indicated. **(D)** Heterotypic interactions between α-PTRAMP Nb 2D11 CDR3 loop (orange) and 3F4 CDR2 loop (green) which both bind to the TSR domain of PTRAMP (light purple). **(E)** α-PTRAMP Nb3F4 nanobody CDR1 – 3 interactions with the TSR domain on PTRAMP. Interacting residues are colored whilst non-interacting residues are white. Hydrogen bonds are denoted as yellow dashed lines. **(F)** α-CSS Nb epitopes on CSS colored in green. Interacting CDRS and FRs are shown.

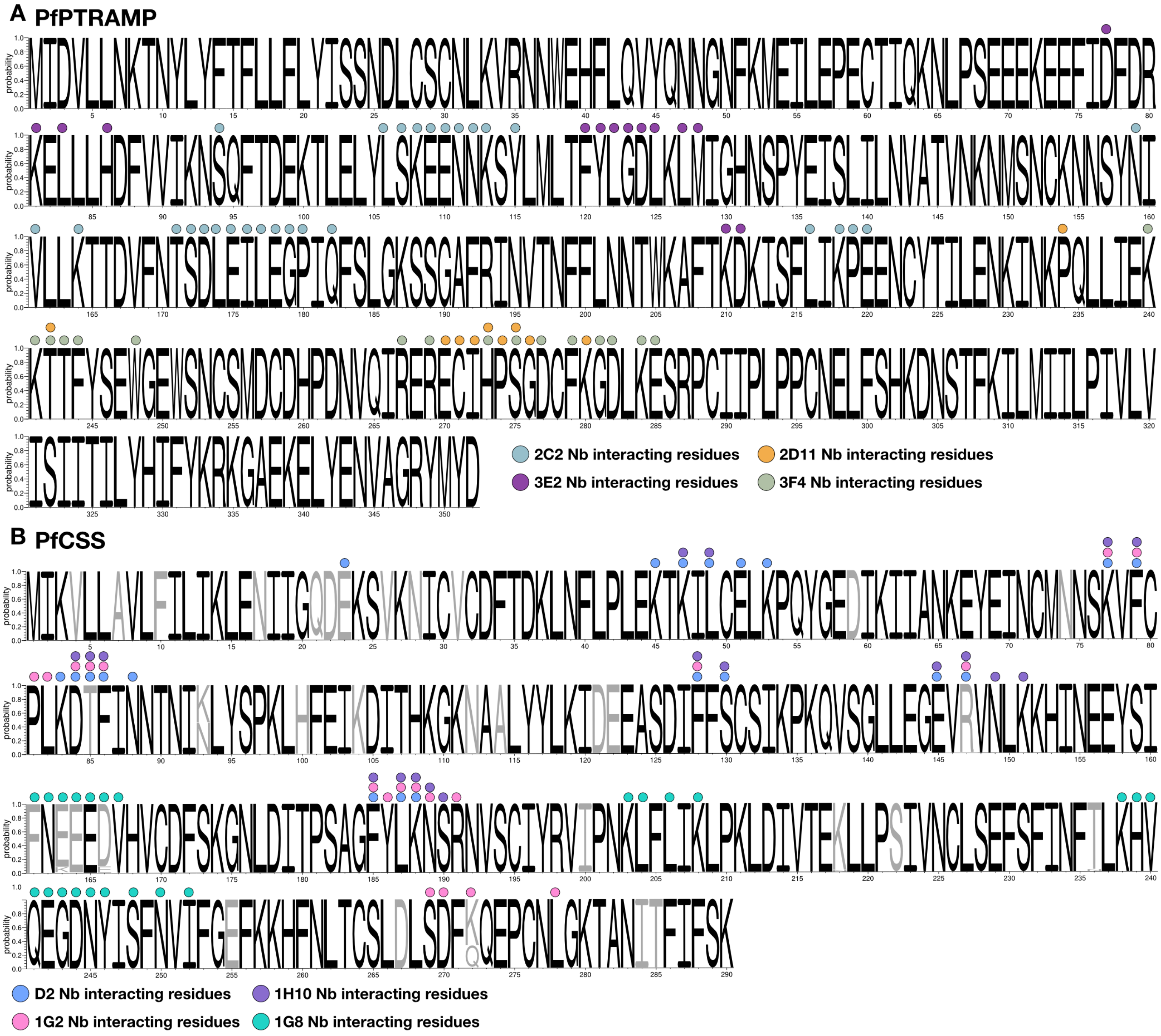


**Supplementary Figure 4. PfPTRAMP-CSS sequence conservation across P. falciparum field isolates (A)** Weblogo representation of PTRAMP sequence diversity among 16,203 *P. falciparum* field isolates from the MalariaGen Pf7 dataset. Residues which interact with indicated nanobodies are represented by a colored circle on top of amino acid residues. **(B)** Weblogo representation of PfCSS sequence diversity from 16,203 *P. falciparum* field samples. Conserved residues are colored in black whilst residues with common mutations are colored in grey. See also Table S14.

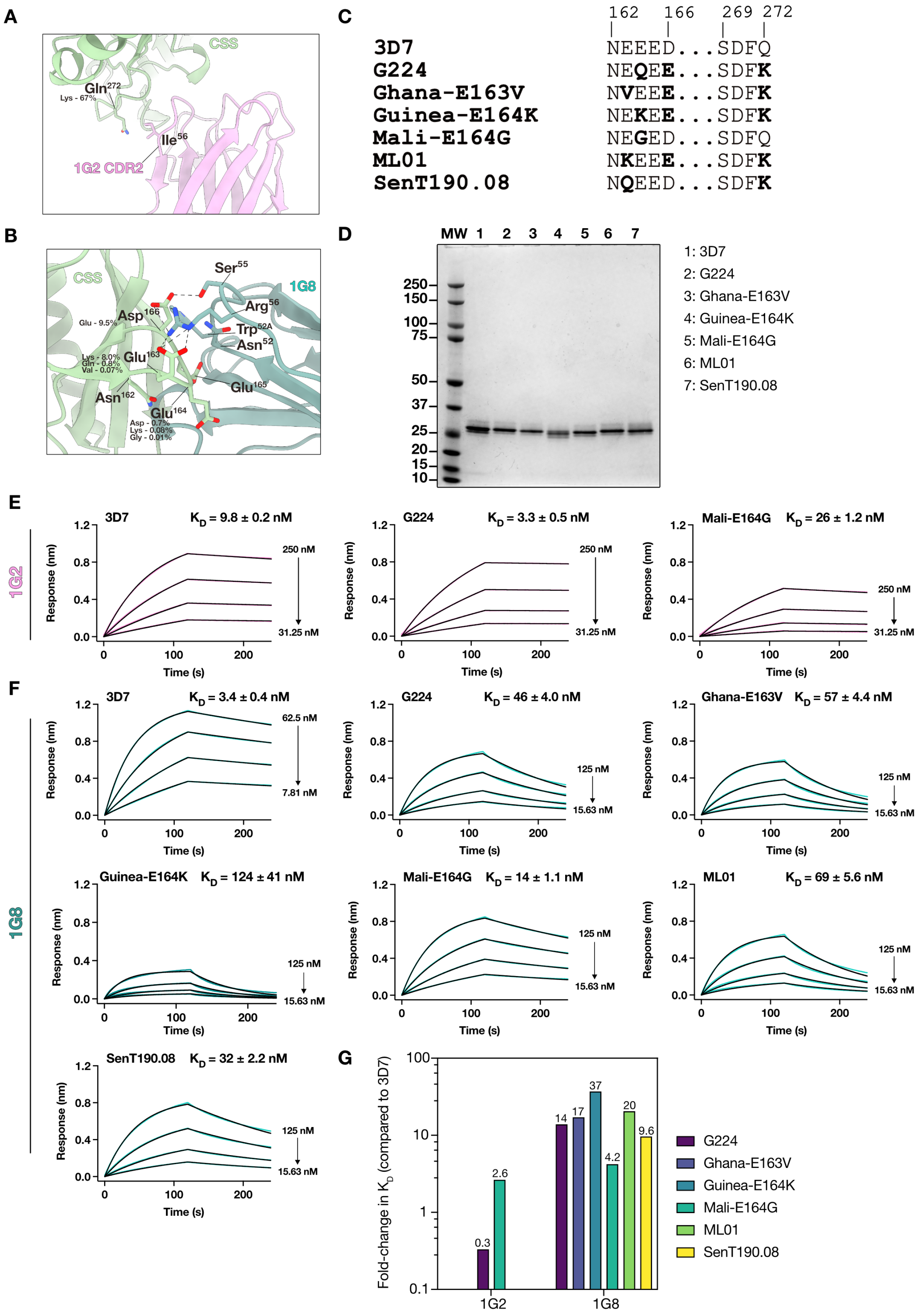


**Supplementary Figure 5. Impact of PfCSS sequence diversity on 1G8 and 1G2 nanobody binding. (A)** Interaction between CDR2 of 1G2 and side chain of Gln 272 on CSS. **(B)** Interaction between 1G8 and residues Asn162 – Asp166 on CSS. Hydrogen bonds are shown by dashed lines. Frequency of mutation at specific residues across global *P. falciparum* isolates from the MalariaGen Pf7 dataset. **(C)** Sequence alignment of haplotypes expressed which contain all possible mutations identified from MalariaGen Pf7 dataset in the 1G8 and 1G2 binding site. Amino acid mutations from 3D7 sequence are shown in bold. **(D)** Coomassie-stained SDS-PAGE gel of different PfCSS variants that were expressed. **(E)** Representative sensorgrams of 1G2 nanobody binding to 3D7, G224 (contains Q272K mutation) and Mali-E164G (does not contain Q272K mutation). **(F)**

Representative sensorgrams of 1G8 nanobody binding to 3D7, G224, Ghana-E163V, Guinea-E164K, Mali-E164G, ML01 and SenT190.08. Dilution series data is shown in pink (1G2) or teal (1G8) whilst 1:1 model of best fit is shown in black. Starting and final concentration of dilution series is also indicated in nM. $K_D$ and SEM (n=2) is also shown. See also Table S15. **(G)** Fold-change in $K_D$ of nanobodies binding to PfCSS mutants compared to 3D7.

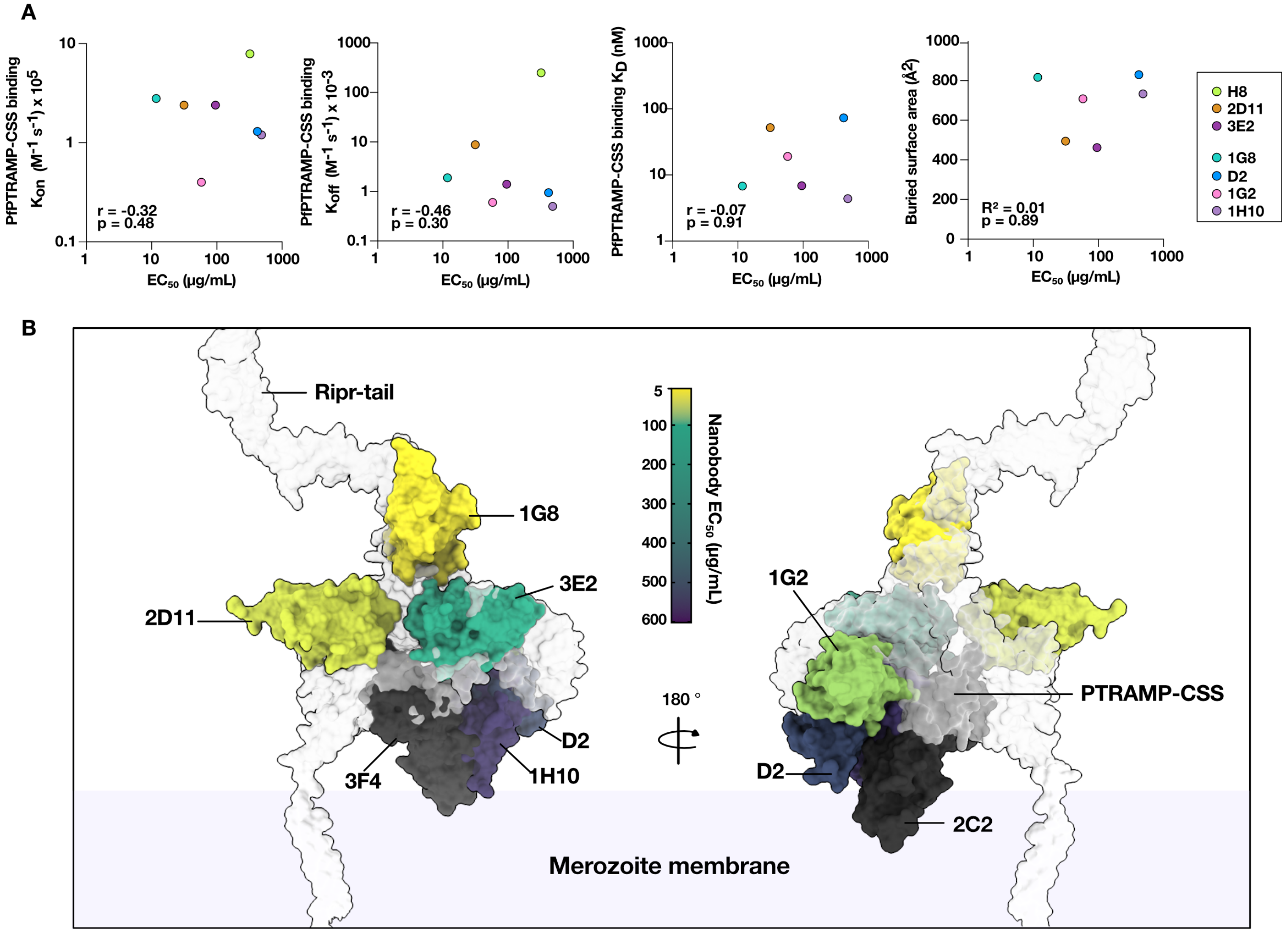


**Supplementary Figure 6. Correlation of nanobody inhibition with kinetic and structural analysis. (A)** PTRAMP-CSS nanobodies binding parameters $K_{on}$, $K_{off}$, and $K_D$ (left to right) correlated with GIA $EC_{50}$ for all inhibitory nanobodies (n = 7). Spearman's rank correlation coefficient (rho values) and two-tailed *p* values are shown. Far right: correlation between GIA $EC_{50}$ and buried surface area between Nb and PTRAMP or CSS using Pearson's correlation, showing r values and *p* values. **(B)** Mapping of non-inhibitory and inhibitory nanobodies onto the AlphaFold3 predicted PTRAMP-CSS-Ripr-tail complex during erythrocyte invasion. Nanobodies are colored based on $EC_{50}$ values, where yellow is the lowest value and purple is the highest. Non-inhibitory nanobodies are shown in black.

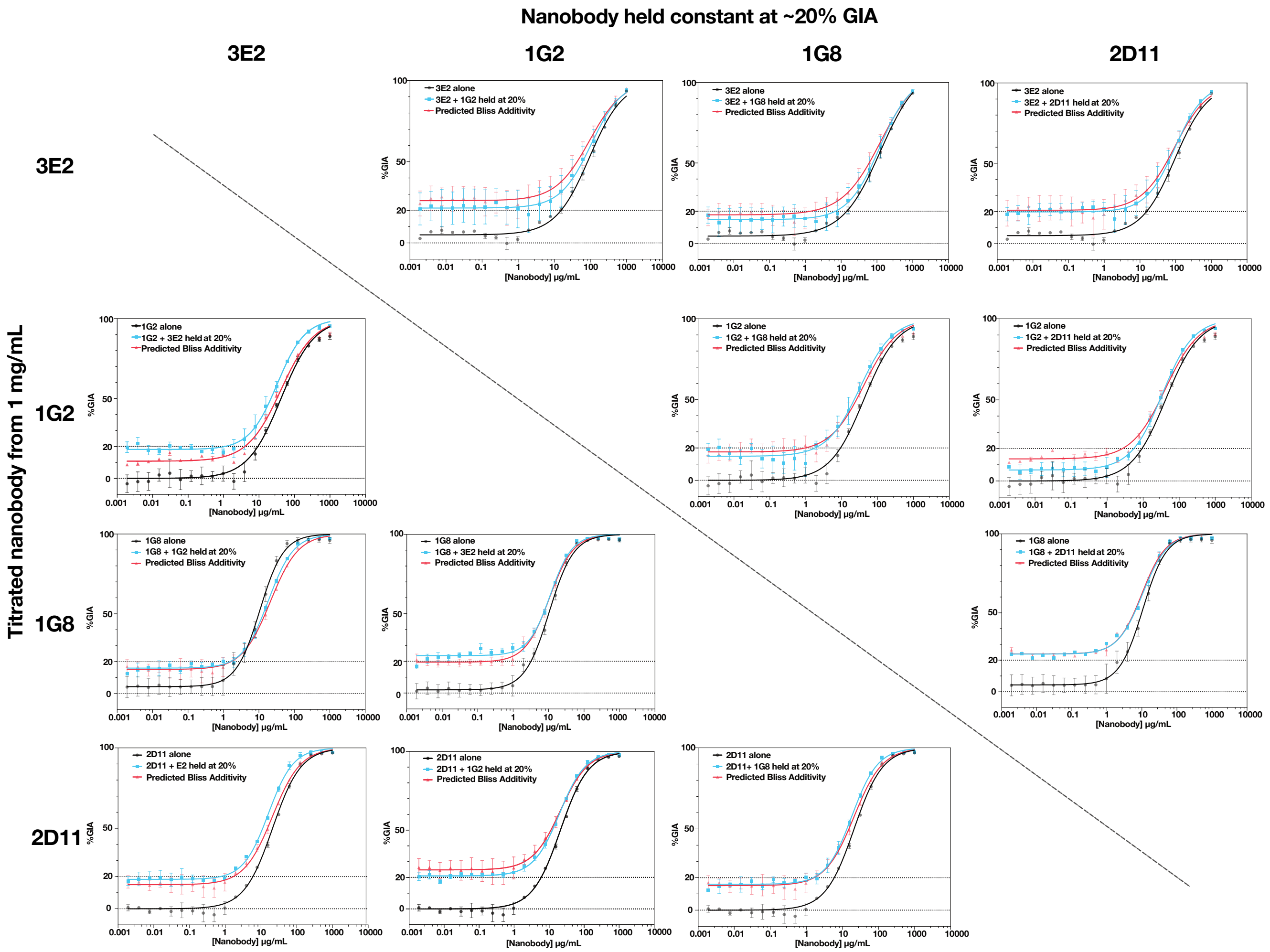


**Supplementary Figure 7. Synergy evaluations between inhibitory nanobodies.** *In vitro* GIA nanobody synergy evaluations with one Nb held at a constant concentration that would yield ~20% GIA and one nanobody is titrated across a 20-step 2-fold dilution series starting at 1 mg/mL (blue). Red indicates the predicted Bliss Additivity. Black indicates the %GIA of the titrated nanobody alone from the same experiment. Each point is the mean of two independent experiments performed in triplicate. Errors bars indicate SEM. Combinations were performed in a pairwise formation for all inhibitory Nbs except for 1H10.

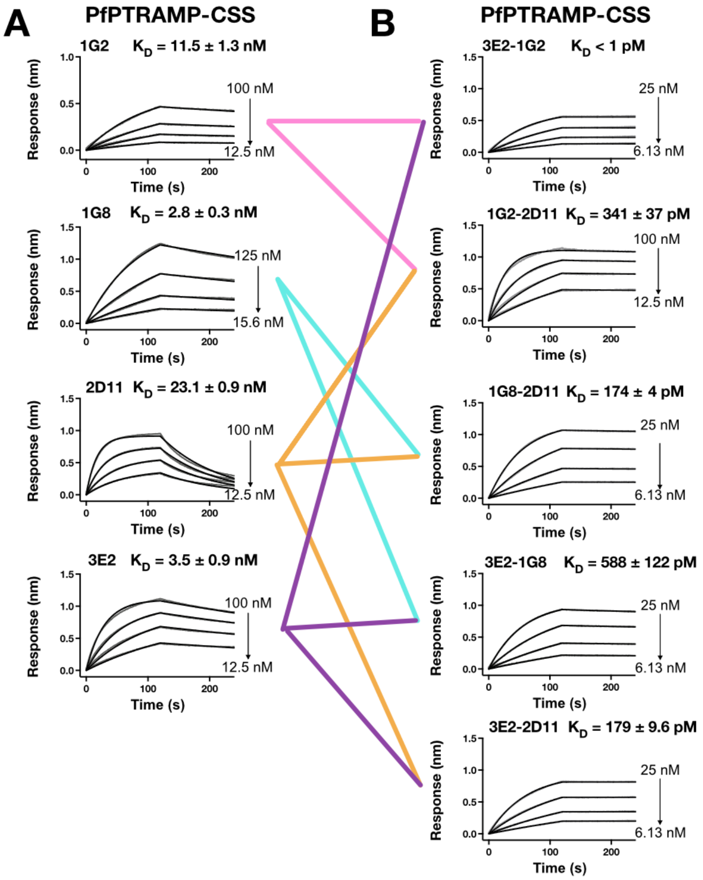


**Supplementary Figure 8. Binding kinetics of nanobody-Fc and bispecific nanobody-Fc proteins to *P. falciparum* PTRAMP-CSS. (A)** Representative sensorgrams of Nb-Fcs binding to PTRAMP-CSS. Dilution series data is shown in grey whilst 1:1 model of best fit is shown in black. Starting and final concentration of dilution series is also indicated in nM. $K_D$ and SEM (n=2) is also shown. See also Table S16. **(B)** Representative sensorgrams of bispecific Nb-Fcs binding to PTRAMP-CSS. Colored lines represent which Nbs (A) are included in BsNb-Fcs tested for binding.

### Table S1. Binding kinetics of nanobodies to PTRAMP, CSS and PTRAMP-CSS.

| | PfPTRAMP | | | PfPTRAMP-CSS | | | |
|---|---|---|---|---|---|---|---|
| **Nb** | **$K_D$ (nM)** | **$k_a$ (1/Ms) x $10^5$** | **$k_d$ (1/s) x $10^{-3}$** | **$K_D$ (nM)** | **$k_a$ (1/Ms) x $10^5$** | **$k_d$ (1/s) x $10^{-3}$** | **Fold $\Delta K_D$ (increase)** |
| 2C2 | 15.0 ± 5.9 | 1.4 ± 0.4 | 1.8 ± 0.1 | 11.0 ± 3.8 | 1.3 ± 0.7 | 1.2 ± 0.2 | 0.7 |
| 2D12 | 39.1 ± 7.3 | 3.8 ± 0.7 | 14.2 ± 0.0 | 31.3 ± 14.8 | 2.8 ± 1.4 | 6.6 ± 0.3 | 0.8 |
| 2D11 | 64.1 ± 25.3 | 3.0 ± 0.9 | 17 ± 1.8 | 52.0 ± 28.4 | 2.4 ± 1.4 | 8.8 ± 0.1 | 0.8 |
| 3E2 | 9.5 ± 1.3 | 3.2 ± 0.1 | 3.0 ± 0.4 | 6.9 ± 2.4 | 2.4 ± 0.9 | 1.4 ± 0.1 | 0.7 |
| 3F4 | 35.2 ± 3.0 | 0.9 ± 0.1 | 3.0 ± 0.2 | 24.1 ± 16.5 | 1.3 ± 0.9 | 1.8 ± 0.1 | 0.7 |
| | **PfCSS** | | | **PfPTRAMP-CSS** | | | |
| **Nb** | **$K_D$ (nM)** | **$k_a$ (1/Ms) x $10^5$** | **$k_d$ (1/s) x $10^{-3}$** | **$K_D$ (nM)** | **$k_a$ (1/Ms) x $10^5$** | **$k_d$ (1/s) x $10^{-3}$** | **Fold $\Delta K_D$ (increase)** |
| 1A7 | 11.4 ± 0.7 | 1.9 ± 0.5 | 2.1 ± 0.4 | 11.3 ± 1.4 | 3.4 ± 1.0 | 3.7 ± 0.7 | 1.0 |
| 1A8 | 3.4 ± 0.1 | 1.7 ± 0.9 | 0.6 ± 0.3 | 4.3 ± 1.2 | 3.4 ± 1.9 | 1.2 ± 0.4 | 1.3 |
| 1B2 | 8.7 ± 1.1 | 1.6 ± 0.6 | 1.3 ± 0.3 | 7.7 ± 1.2 | 3.3 ± 1.1 | 2.5 ± 0.5 | 0.9 |
| 1B7 | 5.7 ± 0.8 | 1.5 ± 0.5 | 0.8 ± 0.2 | 5.9 ± 0.5 | 2.5 ± 0.8 | 1.4 ± 0.3 | 1.0 |
| 1B11 | 1.9 ± 0.2 | 2.0 ± 0.7 | 0.4 ± 0.2 | 2.0 ± 0.2 | 3.4 ± 1.1 | 0.7 ± 0.2 | 1.1 |
| 1C3 | 0.9 ± 0.3 | 1.1 ± 0.4 | 0.1 ± 0.4 | 5.6 ± 2.3 | 1.5 ± 0.2 | 0.9 ± 0.5 | 6.0 |
| 1C6 | 3.2 ± 1.9 | 1.5 ± 0.6 | 0.4 ± 0.1 | 4.7 ± 0.4 | 2.0 ± 0.7 | 0.9 ± 0.3 | 1.5 |
| 1D4 | 7.7 ± 1.3 | 1.5 ± 0.3 | 1.1 ± 0.0 | 6.1 ± 1.1 | 2.7 ± 0.8 | 1.6 ± 0.2 | 0.8 |
| 1E1 | 7.6 ± 3.2 | 0.7 ± 0.0 | 0.5 ± 0.2 | 1.9 ± 1.6 | 1.3 ± 0.5 | 0.2 ± 0.1 | 0.3 |
| 1E3 | 11.9 ± 2.9 | 1.5 ± 0.4 | 1.7 ± 0.0 | 9.2 ± 1.6 | 2.8 ± 0.9 | 2.4 ± 0.4 | 0.8 |
| 1E7 | 13.9 ± 2.4 | 1.1 ± 0.3 | 1.4 ± 1.4 | 19.0 ± 1.8 | 1.6 ± 0.3 | 3.1 ± 0.8 | 1.4 |
| 1F2 | N.B. | N.B. | N.B. | 41.8 ± 0.1 | 6.2 ± 0.1 | 2.6 ± 0.4 | - |
| 1F10 | 10.1 ± 0.5 | 1.7 ± 0.5 | 1.7 ± 0.5 | 8.1 ± 1.3 | 3.2 ± 0.9 | 2.5 ± 0.3 | 0.8 |
| 1G2 | 14.2 ± 2.8 | 0.4 ± 0.2 | 0.5 ± 0.2 | 19.1 ± 13.3 | 0.4 ± 0.2 | 0.6 ± 0.3 | 1.4 |
| 1G8 | 5.9 ± 1.1 | 1.7 ± 0.4 | 1.0 ± 0.4 | 6.8 ± 0.4 | 2.8 ± 0.5 | 1.9 ± 0.4 | 1.2 |
| 1G11 | 6.3 ± 0.7 | 1.5 ± 0.5 | 0.9 ± 0.2 | 5.1 ± 0.6 | 2.5 ± 0.6 | 1.2 ± 0.2 | 0.8 |
| 1G12 | 2.5 ± 0.4 | 1.2 ± 0.5 | 0.3 ± 0.1 | 2.2 ± 0.5 | 2.3 ± 0.7 | 0.5 ± 0.01 | 0.9 |
| 1G1 | 4.4 ± 0.7 | 1.6 ± 0.6 | 0.6 ± 0.2 | 3.8 ± 0.8 | 2.9 ± 0.7 | 1.0 ± 0.1 | 0.9 |
| 1H7 | 6.6 ± 0.8 | 1.6 ± 0.6 | 1.0 ± 0.3 | 6.0 ± 0.2 | 3.0 ± 0.5 | 1.8 ± 0.2 | 0.9 |
| 1H10 | 24.7 ± 2.3 | 1.3 ± 0.5 | 3.2 ± 1.0 | 4.4 ± 0.3 | 1.2 ± 0.2 | 0.5 ± 0.1 | 0.2 |

**Table S2. Binding kinetics of PTRAMP-specific nanobodies to the TSR and Growth Factor Domains of PTRAMP.**

| | TSR | | | Growth Factor Domain | | |
|---|---|---|---|---|---|---|
| **Nb** | **$K_D$ (nM)** | **$k_a$ (1/Ms) x $10^5$** | **$k_d$ (1/s) x $10^{-3}$** | **$K_D$ (nM)** | **$k_a$ (1/Ms) x $10^5$** | **$k_d$ (1/s) x $10^{-3}$** |
| 2C2 | N.B. | | | 17.5 ± 0.7 | 1.16 ± 0.4 | 1.8 ± 0.2 |
| 2D12 | N.B. | | | 38.6 ± 11.8 | 2.3 ± 0.5 | 8.4 ± 0.9 |
| 2D11 | 108 ± 77 | 8.3 ± 3.3 | 64.1 ± 28.3 | N.B. | | |
| 3E2 | N.B. | | | 70.1 ± 20.5 | 2.3 ± 0.5 | 15.2 ± 1.1 |
| 3F4 | 15.3 ± 4.0 | 2.5 ± 0.0 | 3.8 ± 1.0 | N.B. | | |

**Table S3. $EC_{50}$ (µg/mL) of Nbs in asexual-stage *P. falciparum* growth inhibition assays.**

| **Nanobody** | **$EC_{50}$ (µg/mL) (± 95% confidence interval)** | **Fold decrease in $EC_{50}$ from previously characterised nanobody*** |
|---|---|---|
| 1G8 | 11.8 (10.5 – 13.2) | 35.5 |
| 2D11 | 31.4 (26.9 – 36.6) | 10.3 |
| 1G2 | 58.1 (46.4 – 72.5) | 7.2 |
| 3E2 | 95.6 (86.1 – 106) | 3.4 |
| H8 | 322 (283 – 368) | - |
| 1H10 | 485 (379 – 621) | 0.9 |
| D2 | 419 (260 – 736) | - |

*PTRAMP specific Nbs were compared to the $EC_{50}$ of H8 and CSS specific Nbs were compared to D2 to calculate fold-change in $EC_{50}$.

### Table S4: Data collection and refinement statistics for Nb-PTRAMP-CSS complexes

| | 1G2-CSS | 1G8-PTRAMP-CSS | 1H10-CSS | 2C2-PTRAMP | 2D11-3F4-PTRAMP-TSR | 3E2-PTRAMP-GFD |
|---|---|---|---|---|---|---|
| **Beamline** | MX2 | MX2 | MX2 | MX2 | MX2 | MX2 |
| **Wavelength (Å)** | 0.953732 | 0.95373 | 0.95373 | 0.953739 | 0.953737 | 0.953737 |
| **Space group** | $P2_1$ | $P6_522$ | C2 | $P2_1$ | C2 | P1 |
| **Cell dimensions** | | | | | | |
| ***a,b,c* (Å)** | 83.1, 71.7, 94.6 | 153.7, 153.7, 341.2 | 94.3, 96.7, 52.7 | 54.7, 73.4, 55.6 | 151.8, 68.0, 74.1 | 60.6, 64.2, 86.0 |
| **α, β, γ (°)** | 90.0, 108.8, 90.0 | 90.0, 90.0, 120.0 | 90.0, 95.0, 90.0 | 90.0, 111.1, 90.0 | 90.0, 111.6, 90.0 | 76.5, 87.1, 85.0 |
| **Resolution (Å)[a]** | 38.58-3.23 (3.35-3.23) | 4-4.20 (4.60-4.20) | 42.7-2.2 (2.37-2.2) | 40.0-1.50 (1.53-1.5) | 40.0-2.12 (2.18-2.12) | 40.0-2.40 (2.48-2.40) |
| **No. molecules in ASU** | 2 | 2 | 1 | 1 | 2 | 4 |
| **No. observations** | 57,075 (11,976) | 237,228 (55,766) | 50,215 (2,423) | 227,072 (11,083) | 198,641 (17,177) | 88,100 (7,893) |
| **No. unique observations** | 16,981 (1,676) | 18,193 (4,236) | 14,422 (721) | 65,559 (3,249) | 39,314 (3,234) | 47,965 (4,400) |
| **Multiplicity** | 3.4 (3.4) | 13.0 (13.2) | 3.5 (3.4) | 3.5 (3.4) | 9.8 (2.7) | 1.8 (1.8) |
| **$R_{merge}$ (%)** | 4.4 (66.4) | 16.7 (173.50) | 6.2 (47.2) | 6.1 (75.8) | 9.5 (74.9) | 6.4 (66.6) |
| **$R_{pim}$ (%)** | 9.5 (48.4) | 4.7 (49.0) | 5.08 (46.8) | 3.9 (48.2) | 7.3 (48.8) | 8.3 (49.9) |
| **$\langle I/\sigma I\rangle$** | 6.5 (1.6) | 10.5 (1.5) | 11.7 (2.1) | 11.1 (1.7) | 9.8 (2.7) | 6.2 (1.1) |
| **$CC_{1/2}$** | 99.2 (67.5) | 99.9 (79.4) | 99.9 (69.6) | 99.8 (63.0) | 98.6 (46.1) | 99.4 (55.0) |
| **Completeness (%)** | 99.9 (99.5) | 99.9 (98.3) | 91.9 (83.8) | 99.9 (99.9) | 98.6 (99.2) | 97.9 (97.3) |
| **Refinement Statistics** | | | | | | |
| **Reflections used in refinement** | 16, 971 | 18,086 | 14,414 | 65,528 | 39,293 | 47,980 |
| **Reflections used for R-free** | 838 | 1,760 | 1,440 | 3,336 | 2,023 | 2,412 |
| **Non-hydrogen atoms** | 6,319 | 6,630 | 3,188 | 3,309 | 5,067 | 9,154 |
| **Macromolecule** | 6,184 | 6,552 | 3,118 | 2,832 | 4,723 | 9,007 |
| **Water** | 21 | 0 | 28 | 451 | 338 | 123 |
| **Heteroatom** | 114 | 78 | 42 | 26 | 6 | 24 |
| **$R_{work}$ / $R_{free}$** | 20.9 / 26.8 | 15.8 / 18.4 | 22.5 / 28.2 | 16.9 / 18.8 | 20.5 / 24.2 | 22.8 / 25.7 |
| **Rms deviations from ideality** | | | | | | |
| **Bond lengths (Å)** | 0.003 | 0.008 | 0.002 | 0.014 | 0.003 | 0.002 |
| **Bond angle (°)** | 0.65 | 1.17 | 0.48 | 1.27 | 0.56 | 0.47 |
| **Ramachandran plot** | | | | | | |
| **Favoured regions (%)** | 93.1 | 91.0 | 96.4 | 97.4 | 98.3 | 96.5 |
| **Allowed regions (%)** | 6.9 | 9.0 | 3.6 | 2.6 | 1.7 | 3.5 |
| **B-factors ($Å^2$)** | | | | | | |
| **Wilson B-value** | 80.6 | 198.0 | 38.4 | 17.7 | 27.2 | 47.2 |
| **Average B-factors** | 88.9 | 249.5 | 47.8 | 25.8 | 35.0 | 57.4 |
| **Average macromolecule** | 88.5 | 248.4 | 47.7 | 23.8 | 34.9 | 57.5 |
| **Average heteroatom** | 114.2 | 340.2 | 63.1 | 47.2 | 28.9 | 60.1 |
| **Average water molecule** | 54.6 | - | 38.6 | 36.5 | 36.3 | 51.1 |

[a] Values in parentheses refer to the highest resolution bin

**Table S5. Table of contacts between PTRAMP and 2C2.**

| PTRAMP Residue (BSA Å$^2$) | Interaction Type | 2C2 Residue |
|---|---|---|
| **Ser94 (40.7)** | | |
| Ser | VDW | Leu96, Tyr100H, Asp101, Phe102 |
| Ser$^{OG}$ | HB | Asp101$^{O}$, Asp101$^{OD1}$ |
| **Leu106 (27.0)** | | |
| Leu | VDW | Tyr100B, Asn100F, Ala100G |
| **Ser107 (12.5)** | | |
| Ser | VDW | Asn100F, Ala100G |
| **Lys108 (58.9)** | | |
| Lys | VDW | Asp100D, Asn100F |
| Lys$^{O}$ | HB | Asn100F$^{ND2}$ |
| **Glu109 (4.5)** | | |
| Glu | VDW | Asn100F |
| **Glu110 (9.2)** | | |
| Glu | VDW | Asn100F |
| **Asn111 (40.0)** | | |
| Asn | VDW | Asn100F, Trp103 |
| **Asn112 (0.3)** | | |
| Asn | VDW | Tyr100H |
| Asn$^{N}$ | WMHB | Asn100F$^{O}$ |
| **Lys113 (42.3)** | | |
| Lys | VDW | Asn100F, Ala100G, Tyr100H, Asp101, Trp103 |
| Lys$^{NZ}$ | HB | Asn100F$^{O}$, Tyr100H$^{O}$ |
| **Tyr115 (15.2)** | | |
| Tyr | VDW | Leu96, Ala100G, Tyr100H, Asp101 |
| Tyr$^{OH}$ | HB | Asp101$^{OD1}$ |
| **Asn159 (33.0)** | | |
| Asn | VDW | Ala97, Gly98, Arg100A, Tyr100B |
| Asn$^{ND2}$ | HB | Tyr100B$^{OH}$ |
| **Val161 (12.1)** | | |
| Val | VDW | Leu96, Ala97, Gly98 |
| **Lys164 (8.8)** | | |
| Lys | VDW | Arg27 |
| **Thr171 (23.7)** | | |
| Thr | VDW | Tyr32, Leu96, Phe102 |
| Thr$^{O}$ | HB | Tyr32$^{OH}$ |
| **Ser172 (95.5)** | | |
| Ser | VDW | Gln3, Ala28, Phe29, Tyr32, Phe102 |
| SerN | WMHB | Ala$^{N}$ |
| Ser$^{OG}$ | HB | Gln3$^{NE2}$ |
| **Asp173 (3.3)** | | |
| Asp | VDW | Ala28, Tyr32 |
| **Leu174 (28.8)** | | |
| Leu | VDW | Ala28, Thr31, Tyr32 |
| Leu$^{N}$ | HB | Tyr32$^{OH}$ |
| Leu$^{O}$ | HB | Tyr32$^{OH}$ |
| **Glu175 (67.4)** | | |
| Glu | VDW | Arg27, Ala28, Thr31 |
| Glu$^{OE2}$ | HB | Arg27$^{NH1}$ |
| Glu$^{OE1}$ | SB | Arg27$^{NH1}$ |
| Glu$^{OE2}$ | SB | Arg27$^{NH1}$ |
| **Ile176 (88.2)** | | |
| Ile | VDW | Thr31, Tyr32, Asp95, Leu96, Ala97, Gly98 |
| Ile$^{N}$ | HB | Thr31$^{OG1}$ |
| **Leu177 (16.8)** | | |
| Leu | VDW | Arg53 |
| Leu$^{O}$ | HB | Arg53$^{NH2}$ |

| | | |
|---|---|---|
| **Glu178 (24.4)** | | |
| Glu | VDW | Arg53 |
| $Glu^{OE1}$ | HB | $Arg53^{NH1}$, $Arg53^{NH2}$ |
| $Glu^{OE1}$ | SB | $Arg53^{NH1}$, $Arg53^{NH2}$ |
| $Glu^{OE2}$ | SB | $Arg53^{NH2}$ |
| **Gly179 (10.1)** | | |
| Gly | VDW | Ala97, Gly98 |
| **Pro180 (43.3)** | | |
| Pro | VDW | Ala97, Gly98, Ser99 |
| **Gln182 (31.8)** | | |
| Gln | VDW | Arg100A |
| **Leu216 (27.9)** | | |
| Leu | VDW | Leu96 |
| **Lys218 (82.2)** | | |
| Lys | VDW | Asp95, Ala97, Tyr100B, Ala100G, Tyr100H, Asp101 |
| $Lys^{NZ}$ | HB | $Asp95^{OD1}$, $Ala100G^{O}$, $Asp101^{OD2}$ |
| $Lys^{NZ}$ | SB | $Asp95^{OD1}$, $Asp101^{OD2}$ |
| $Lys^{O}$ | HB | $Tyr100B^{OH}$ |
| **Pro219 (2.0)** | | |
| Pro | VDW | Tyr100B |
| $Pro^{O}$ | HB | $Tyr100B^{OH}$ |
| **Glu220 (49.3)** | | |
| Glu | VDW | Arg100A, Tyr100B |
| $Glu^{OE1}$ | HB | $Arg100A^{NH2}$ |
| $Glu^{OE2}$ | HB | $Arg100A^{NE}$ |
| $Glu^{OE1}$ | SB | $Arg100A^{NE}$, $Arg100A^{NH2}$ |
| $Glu^{OE2}$ | SB | $Arg100A^{NE}$, $Arg100A^{NH2}$ |

Van Der Waals (VDW)
Hydrogen Bond (HB)
Salt Bridge (SB)
Water-mediated hydrogen bond (WMHB)

### Table S6. Table of contacts between PTRAMP and 3E2.

| PTRAMP Residue (BSA Å$^2$) | Interaction Type | 3E2 Residue |
|---|---|---|
| **Asp77 (19.7)** | | |
| Asp | VDW | Arg53 |
| $Asp^{OD1}$ | HB | $Arg53^{NH2}$ |
| $Asp^{OD1}$ | SB | $Arg53^{NH2}$ |
| **Lys81 (32.0)** | | |
| Lys | VDW | Asp96, Ile97 |
| $Lys^{NZ}$ | HB | $Asp96^{OD1}$ |
| $Lys^{NZ}$ | SB | $Asp96^{OD1}$ |
| **Leu83 (9.7)** | | |
| Leu | VDW | Ile97 |
| **His86 (10.5)** | | |
| His | VDW | Ser100C |
| **Tyr121 (86.9)** | | |
| Tyr | VDW | Ile97, Gly98, Gly99, Ala100, Ser100A, Ala100D |
| $Tyr^{OH}$ | HB | $Gly98^{O}$, $Ala100^{N}$, $Ser100A^{N}$ |
| **Leu122 (3.9)** | | |
| Leu | VDW | Ile97 |
| **Gly123 (9.7)** | | |
| Gly | VDW | Ile97, Gly98, Gly99 |
| **Asp124 (108.3)** | | |
| Asp | VDW | Ile33, Arg50, Thr52, Phe58, Ile97, Gly98, |

| | | |
|---|---|---|
| | | Gly99, Tyr100E |
| $Asp^{N}$ | HB | $Ile97^{O}$ |
| $Asp^{OD1}$ | HB | $Arg50^{NH2}$ |
| $Asp^{OD2}$ | HB | $Gly99^{N}$, $Arg50^{NE}$ |
| $Asp^{OD1}$ | SB | $Arg50^{NH2}$, $Arg50^{NE}$ |
| $Asp^{OD2}$ | SB | $Arg50^{NH2}$, $Arg50^{NE}$ |
| **Leu125 (40.1)** | | |
| Leu | VDW | Thr52, Asp96, Ile97 |
| $Leu^{N}$ | HB | $Ile97^{O}$ |
| **Leu127 (71.1)** | | |
| Leu | VDW | Thr52, Tyr56, Phe58 |
| **Met128 (19.1)** | | |
| Met | VDW | Thr52 |
| **Lys210 (20.4)** | | |
| Lys | VDW | Ala100 |
| **Asp211 (30.3)** | | |
| Asp | VDW | Arg50, Gly99, Ala100, Ser100A |
| $Asp^{OD2}$ | HB | $Ala100^{N}$ |
| $Asp^{OD1}$ | SB | $Arg50^{NH2}$ |

Van Der Waals (VDW)
Hydrogen Bond (HB)
Salt Bridge (SB)
Water-mediated hydrogen bond (WMHB)

### Table S7. Table of contacts between PTRAMP and 2D11.

| PTRAMP Residue (BSA Å$^2$) | Interaction Type | 2D11 Residue |
|---|---|---|
| **Pro241 (16.4)** | | |
| Pro | VDW | Ser99 |
| **Thr243 (10.3)** | | |
| Thr | VDW | Ser99 |
| **Glu270 (55.9)** | | |
| Glu | VDW | Ser100A, Tyr100B, Tyr100C |
| $Glu^{OE2}$ | HB | $Ser100A^{OG}$ |
| **Cys271 (31.1)** | | |
| Cys | VDW | Tyr58, Gly100, Tyr100B |
| $Cys^{SG}$ | HB | $Tyr100B^{OH}$ |
| **Ile272 (62.0)** | | |
| Ile | VDW | Lys95, Val98, Ser99, Gly100, Tyr100B |
| $Ile^{O}$ | HB | $Gly100^{N}$ |
| **His273 (6.0)** | | |
| His | VDW | Ser99, Tyr100B |
| $His^{N}$ | HB | $Tyr100B^{OH}$ |
| $His^{O}$ | HB | $Tyr100B^{OH}$ |
| **Pro274 (104.5)** | | |
| Pro | VDW | Asn31, Leu32, Leu33, Lys95, Val98, Lys95, Ser96, Pro97, Val98, Ser99, Tyr100B |
| $Pro^{O}$ | HB | $Leu33^{N}$ |
| **Ser275 (56.5)** | | |
| Ser | VDW | Asn31, Leu32, Leu33, Ile51, Ser52, Trp52A |
| $Ser^{O}$ | HB | $Trp52A^{N}$ |
| **Gly276 (58.2)** | | |
| Gly | VDW | Leu33, Ser52, Ser56, Tyr58, Tyr100B |
| $Gly^{O}$ | HB | $Tyr58^{OH}$ |
| **Asp277 (57.8)** | | |
| Asp | VDW | Ser52, Trp52A, Ser53, Gly54, Asp55, Ser56, Tyr58, Tyr100B, |

| | | |
|---|---|---|
| Asp$^{OD1}$ | HB | Ser52$^{OG}$, Ser53$^{OG}$, Ser56$^{N}$ |
| Asp$^{OD2}$ | HB | Ser53$^{N}$, Ser53$^{OG}$ |
| **Cys278 (8.2)** | | |
| Cys | VDW | Ser56, Tyr58 |
| Cys$^{O}$ | HB | Ser56$^{OG}$ |
| **Lys280 (28.0)** | | |
| Lys | VDW | Asp55, Ser56 |

Van Der Waals (VDW)
Hydrogen Bond (HB)

Salt Bridge (SB)
Water-mediated hydrogen bond (WMHB)

### Table S8. Table of contacts between PTRAMP and 3F4.

| PTRAMP Residue (BSA Å$^2$) | Interaction Type | 3F4 Residue |
|---|---|---|
| **Asp240 (27.7)** | | |
| Asp | VDW | Ile100C |
| **Pro241 (10.7)** | | |
| Pro | VDW | Tyr32, Ile100C |
| **Thr242 (26.2)** | | |
| Thr | VDW | Asn31, Tyr32 |
| Thr$^{O}$ | HB | Tyr32$^{OH}$ |
| **Thr243 (24.1)** | | |
| Thr | VDW | Thr28, Asn31 |
| **Phe244 (85.8)** | | |
| Phe | VDW | Asn31, Tyr32, Gly96, Ala97, Trp100A, Ile100C |
| Phe$^{N}$ | HB | Asn31$^{OD1}$ |
| **Tyr245 (47.7)** | | |
| Tyr | VDW | Asp30, Asn31, Thr52A, Ala97, Trp100A |
| Tyr$^{N}$ | HB | Asn31$^{OD1}$ |
| Tyr$^{OH}$ | HB | Ala97$^{O}$ |
| Tyr$^{OH}$ | WMHB | Ala97$^{N}$ |
| **Trp248 (25.4)** | | |
| Trp | VDW | Asp30, Val53 |
| Trp$^{NE1}$ | HB | Asp30$^{OD2}$ |
| **Arg267 (2.5)** | | |
| Arg | VDW | Asp30 |
| **Arg269 (46.3)** | | |
| Arg | VDW | Asp30, Asn31, Thr52A, Val53 |
| Arg$^{NH2}$ | HB | Asp30$^{O}$ |
| Arg$^{NH1}$ | SB | Asp30$^{OD1}$, Asp30$^{OD2}$ |
| **His273 (38.3)** | | |
| His | VDW | Trp100A, Ile100C |
| **Ser275 (11.5)** | | |
| Ser | VDW | Ser100, Trp100A |
| **Asp277 (53.5)** | | |
| Asp | VDW | Gly98, Ser99, Trp100A |
| Asp$^{OD2}$ | HB | Ser100$^{OG}$ |
| Asp$^{O}$ | HB | Trp100A$^{NE1}$ |
| **Cys278 (13.5)** | | |
| Cys | VDW | Gly98, Ser99, Trp100A |
| **Phe279 (92.0)** | | |
| Phe | VDW | Trp33, Ala97, Gly98, Ser99, Arg100B |
| **Gly281 (34.5)** | | |
| Gly | VDW | Asn52, Val53, Ser55 |
| Gly$^{O}$ | HB | Asn52$^{ND2}$, Ser55$^{OG}$ |
| **Asp282 (53.1)** | | |
| Asp | VDW | Trp33, Asn52, Thr52A, Val53, Ala97 |

| | | |
|---|---|---|
| Asp$^{OD1}$ | HB | Thr52A$^{OG1}$, Thr52A$^{N}$ |
| Asp$^{OD2}$ | HB | Trp33$^{NE1}$ |
| Asp$^{O}$ | HB | Thr52A$^{OG1}$ |
| **Lys284 (10.4)** | | |
| Lys | VDW | Val53 |
| **Glu285 (36.2)** | | |
| Glu | VDW | Val53, Asn73 |

Van Der Waals (VDW)
Hydrogen Bond (HB)
Salt Bridge (SB)
Water-mediated hydrogen bond (WMHB)

### Table S9. Table of contacts between 2D11 and 3F4.

| 2D11 Residue (BSA Å$^2$) | Interaction Type | 3F4 Residue |
|---|---|---|
| **Trp52A (19.7)** | | |
| Trp | VDW | Ser100 |
| **Ser53 (19.5)** | | |
| Ser | VDW | Ser99, Ser100 |
| Ser$^{OG}$ | HB | Ser99$^{OG}$ |
| **Asp55 (26.3)** | | |
| Asp | VDW | Ser99 |
| Asp$^{OD2}$ | HB | Ser99$^{OG}$ |

Van Der Waals (VDW)
Hydrogen Bond (HB)

### Table S10. Table of contacts between CSS and 1H10.

| CSS Residue (BSA Å$^2$) | Interaction Type | 1H10 Residue |
|---|---|---|
| **Lys47 (35.3)** | | |
| Lys | VDW | Tyr100E |
| **Leu49 (29.1)** | | |
| Leu | VDW | Tyr100E, Ser100G |
| **Glu51 (51.7)** | | |
| Glu | VDW | Ser100G, Ser100I, Glu100J |
| Glu$^{OE1}$ | HB | Ser100I$^{OG}$ |
| Glu$^{OE2}$ | HB | Ser100G$^{OG}$ |
| **Lys77 (33.3)** | | |
| Lys | VDW | Ser100C, Trp100D |
| Lys$^{NZ}$ | HB | Ser100C$^{O}$ |
| **Phe79 (13.9)** | | |
| Phe | VDW | Ser96, Gly98, Leu99 |
| **Asp84 (55.6)** | | |
| Asp | VDW | Ser96, Asp97, Gly98 |
| Asp$^{OD1}$ | HB | Gly98$^{N}$ |
| **Thr85 (7.7)** | | |
| Thr | VDW | Gly98 |
| **Phe86 (63.8)** | | |
| Phe | VDW | Gly98, Leu99, Trp100D |
| **Phe128 (24.7)** | | |
| Phe | VDW | Leu99, Trp100D |
| **Ser130 (5.6)** | | |
| Ser | VDW | Trp100D |
| **Glu145 (3.4)** | | |
| Glu | VDW | Trp100D |
| **Arg147 (75.5)** | | |

| | | |
|---|---|---|
| Arg | VDW | Trp100D, Tyr100E, Pro100F, Ser100G, Glu100J |
| Arg$^{NH2}$ | HB | Tyr100E$^{O}$, Glu100J$^{OE2}$ |
| Arg$^{NH2}$ | SB | Glu100J$^{OE2}$ |
| **Asn149 (13.4)** | | |
| Asn | VDW | Ser100I |
| **Lys151 (3.2)** | | |
| Lys | VDW | |
| **Phe185 (52.1)** | | |
| Phe | VDW | Thr95, Ser96, Leu99, Glu100J |
| **Leu187 (102.2)** | | |
| Leu | VDW | Tyr32, Ala94, Thr95, Ser96, Asp101, Tyr102 |
| **Lys188 (69.4)** | | |
| Lys | VDW | Thr28, Tyr31, Tyr32, Tyr102 |
| **Asn189 (72.4)** | | |
| Asn | VDW | Gln1, Phe27, Tyr32, Tyr102 |
| Asn$^{N}$ | HB | Tyr32$^{OH}$ |
| Asn$^{OD1}$ | HB | Tyr32$^{OH}$ |
| **Ser190 (20.9)** | | |
| Ser | VDW | Asp101, Tyr102 |
| Ser$^{N}$ | HB | Tyr102$^{OH}$ |
| Ser$^{OG}$ | HB | Tyr102$^{OH}$ |

Van Der Waals (VDW)
Hydrogen Bond (HB)
Salt Bridge (SB)

### Table S11. Table of contacts between CSS and 1G2

| CSS Residue (BSA Å$^{2}$) | Interaction Type | 1G2 Residue |
|---|---|---|
| **Lys77 (12.4)** | | |
| Lys | VDW | Asp99 |
| **Phe79 (14.2)** | | |
| Phe | VDW | Leu98 |
| **Pro81 (17.2)** | | |
| Pro | VDW | Ile52, Ala53 |
| **Leu82 (50.1)** | | |
| Leu | VDW | Ile52, Ile56, Tyr58 |
| Leu82$^{O}$ | HB | Tyr58$^{OH}$ |
| **Asp84 (114.1)** | | |
| Asp | VDW | Thr33, Ile52, Arg95, Ile100B |
| Asp84$^{O}$ | HB | Arg95$^{NH2}$, |
| Asp84$^{OD1}$ | HB, SB | Arg52A$^{NH2}$, Arg52A$^{NE}$ |
| Asp84$^{OD2}$ | HB | Thr33$^{OG1}$, Arg52A$^{NH2}$, Arg52A$^{NE}$, Arg95$^{NE}$ |
| Asp84$^{OD2}$ | SB | Arg52A$^{NH2}$, Arg52A$^{NE}$, Arg95$^{NE}$ |
| **Thr85 (49.07)** | | |
| Thr85 | VDW | Arg95, Asp99, Gly100 |
| Thr85$^{OG1}$ | HB | Leu98$^{O}$, Gly100$^{N}$ |
| **Phe86 (29.92)** | | |
| Phe86 | VDW | Leu98, Asp99 |
| Phe86$^{N}$ | HB | Leu98$^{O}$ |
| **Phe128 (7.81)** | | |
| Phe | VDW | Leu98 |
| **Arg147 (19.0)** | | |
| Arg | VDW | Leu98 (N) |
| **Phe185 (27.4)** | | |

| | | |
|---|---|---|
| Phe | VDW | Arg52A, Leu98 |
| **Tyr186 (8.21)** | | |
| Tyr | VDW | Ala53 |
| **Leu187 (80.1)** | | |
| Leu | VDW | Thr31, Arg52A, Ala53 |
| **Lys188 (107.6)** | | |
| Lys | VDW | Arg52A, Ala53, Ala54, Asp55, Arg71, Asn73 |
| $Lys188^{N}$ | HB | $Arg52A^{O}$ |
| $Lys188^{NZ}$ | HB | $Asn73^{OD1}$, |
| $Lys188^{NZ}$ | SB | $Asp55^{OD1}$, $Asp55^{OD2}$ |
| **Asn189 (47.0)** | | |
| $Asn189^{OG1}$ | HB | $Asn30^{ND2}$ |
| **Arg191 (13.8)** | | |
| $Arg191^{NE}$ | HB | $Ala53^{O}$ |
| **Ser269 (9.5)** | | |
| Ser | VDW | Ala54 |
| **Asp270 (40.8)** | | |
| Asp | VDW | Asp55, Ile56 |
| **Gln272 (30.4)** | | |
| Gln | VDW | Ile56 |
| **Leu278 (29.6)** | | |
| Leu | VDW | Ala54, Ile56 |

Van Der Waals (VDW)
Hydrogen Bond (HB)
Salt Bridge (SB)
Water-mediated hydrogen bond (WMHB)

### Table S12. Table of contacts between PTRAMP-CSS and 1G8.

| CSS Residue (BSA Å$^2$) | Interaction Type | 1G8 Residue |
|---|---|---|
| **Phe161 (5.5)** | | |
| Phe | VDW | Arg56 |
| **Asn162 (29.9)** | | |
| Asn | VDW | Trp52A, Arg56, Leu100 |
| $Asn^{O}$ | HB | $Arg56^{NH2}$ |
| **Glu163 (35.5)** | | |
| Glu | VDW | Arg56, Leu100 |
| $Glu^{OE2}$ | HB | $Arg56^{NE}$ |
| $Glu^{OE1}$ | SB | $Arg56^{NE}$, $Arg56^{NH2}$ |
| $Glu^{OE2}$ | SB | $Arg56^{NE}$ |
| $Glu^{O}$ | HB | $Arg56^{NH2}$ |
| **Glu164 (73.8)** | | |
| Glu | VDW | Tyr32, Asn52, Arg56, Arg95, Leu97, Leu100, Gly100A |
| **Glu165 (116.5)** | | |
| Glu | VDW | Ser33, Ile51, Asn52, Trp52A, Leu96, Leu97, Gly100A |
| $Glu^{OE1}$ | HB | $Trp52A^{N}$ |
| **Asp166 (55.5)** | | |
| Asp | VDW | Asn52, Trp52A, Ser53, Ser55, Arg56 |
| $Asp^{OD2}$ | HB | $Ser55^{OG}$ |
| $Asp^{OD1}$ | SB | $Arg56^{NH1}$, $Arg56^{NH2}$ |
| $Asp^{OD2}$ | SB | $Arg56^{NH1}$ |

| | | |
|---|---|---|
| **Val167 (7.0)** | | |
| Val | VDW | Trp52A, Leu97 |
| **Lys203 (21.3)** | | |
| Lys | VDW | Trp52A, Ser53, Ser55 |
| **Leu204 (41.1)** | | |
| Leu | VDW | Trp52A, Leu97 |
| **Leu206 (25.1)** | | |
| Leu | VDW | Leu97, Ser98 |
| **Lys208 (33.1)** | | |
| Lys | VDW | Ser98, Lys99, Leu100 |
| **Lys238 (47.3)** | | |
| Lys | VDW | Asn73 |
| $Lys^{NZ}$ | HB | $Asn73^{OD1}$ |
| **His239 (77.9)** | | |
| His | VDW | Thr28, Phe29, Ser30, Gly31, Trp52A, Leu97 |
| $His^{NE2}$ | HB | $Phe29^{O}$, $Ser30^{O}$ |
| **Val240 (6.9)** | | |
| Val | VDW | Thr28 |
| **Gln241 (80.2)** | | |
| Gln | VDW | Thr28, Gly31, Tyr32, Leu96, Leu97, Ser98 |
| $Gln^{NE2}$ | HB | $Ser98^{OG}$ |
| **Glu242 (13.5)** | | |
| Glu | VDW | Leu96 |
| **Gly243 (12.5)** | | |
| Gly | VDW | Leu96, Ser98, Lys99 |
| **Asp244 (58.1)** | | |
| Asp | VDW | Lys99, Tyr100D |
| $Asp^{OD1}$ | HB | $Tyr100D^{OH}$ |
| $Asp^{OD2}$ | HB | $Lys99^{NZ}$ |
| $Asp^{OD2}$ | SB | $Lys99^{NZ}$ |
| **Asn245 (6.6)** | | |
| Asn | VDW | Lys99 |
| **Tyr246 (50.4)** | | |
| Tyr | VDW | Leu96, Ser98, Lys99 |
| $Tyr^{OH}$ | HB | $Ser98^{OG}$, $Lys99^{N}$ |
| **Ser248 (1.9)** | | |
| Ser | VDW | Ser98 |
| **Asn250 (9.0)** | | |
| Asn | VDW | Gly31, Leu97 |
| **Ile252 (8.0)** | | |
| Ile | VDW | Trp52A |

Van Der Waals (VDW)
Hydrogen Bond (HB)
Salt Bridge (SB)

### Table S13. BSA (Å$^2$) and contact summary for Nb-PTRAMP-CSS crystal structures

| Nanobody | H-bonds / Salt Bridge | BSA (Å$^2$) | $K_D$ (nM) (n=2) |
|---|---|---|---|
| **2C2** | 25 / 11 | 899.2 | 11.0 |
| **3E2** | 11 / 7 | 461.7 | 6.9 |
| **2D11** | 14 / 0 | 494.9 | 52.0 |
| **3F4** | 15 / 2 | 639.4 | 24.1 |
| **1G2** | 10 / 3 | 708.2 | 19.1 |
| **1H10** | 10 / 1 | 733.2 | 4.5 |
| **1G8** | 13 / 7 | 816.6 | 6.8 |

**Table S14. Frequency of single amino acid mutations in PfCSS from field isolate samples of *P. falciparum* compared to reference 3D7 strain genome**

| Mutation from 3D7 reference genome | Global Frequency Mutation (%) |
|---|---|
| Q272K | 67.033 |
| K93N | 45.816 |
| D166E | 9.513 |
| E163K | 7.971 |
| T236N | 3.318 |
| K218N | 2.795 |
| Q21K | 0.947 |
| E163Q | 0.83 |
| R147H | 0.712 |
| E164Q | 0.658 |
| Q21E | 0.415 |
| I200N | 0.406 |
| S222N | 0.388 |
| D121H | 0.252 |
| V26A | 0.189 |
| D22N | 0.126 |
| K104N | 0.108 |
| E122K | 0.108 |
| H100D | 0.108 |
| D59G | 0.099 |
| E164K | 0.081 |
| E163V | 0.072 |
| N74T | 0.072 |
| N17T | 0.063 |
| E255D | 0.054 |
| F161Y | 0.027 |
| F10L | 0.018 |
| T285I | 0.018 |
| T85A | 0.018 |
| K93Q | 0.018 |
| N112H | 0.018 |
| K218E | 0.018 |
| D267N | 0.009 |
| V26I | 0.009 |
| I284L | 0.009 |
| E23K | 0.009 |
| V4I | 0.009 |
| V31I | 0.009 |
| E164G | 0.009 |
| A7T | 0.009 |

**Table S15. Binding kinetics of 1G2 and 1G8 Nbs to different PfCSS haplotypes containing mutations in the nanobody binding site.**

| **1G2 Nanobody** | | | | |
|---|---|---|---|---|
| **Isolate name** | $K_D$ (nM) | **$k_a$ (1/Ms) x $10^5$** | **$k_d$ (1/s) x $10^{-3}$** | **Fold $\Delta K_D$ (isolate/3D7)** |
| 3D7[a] | 9.8 ± 0.2 | 0.55 ± 0.02 | 0.54 ± 0.01 | - |
| G224[a] | 3.3 ± 0.5 | 0.35 ± 0.003 | 0.12 ± 0.02 | 0.33 |
| Mali-E164G[b] | 26 ± 1.2 | 0.31 ± 0.003 | 0.79 ± 0.04 | 2.63 |
| **1G8 Nanobody** | | | | |
| **Isolate name** | **$K_D$ (nM)** | **$k_a$ (1/Ms) x $10^5$** | **$k_d$ (1/s) x $10^{-3}$** | **Fold $\Delta K_D$ (isolate/3D7)** |
| 3D7 | 3.4 ± 0.36 | 3.1 ± 0.08 | 1.1 ± 0.1 | - |
| G224 | 46 ± 4.0 | 1.6 ± 0.01 | 7.2 ± 0.7 | 13.8 |
| Ghana-E163V[b] | 57 ± 4.4 | 1.7 ± 0.005 | 9.6 ± 0.8 | 17.02 |
| Guinea-E164K[b] | 124 ± 41 | 1.7 ± 0.27 | 19 ± 3.3 | 36.8 |
| Mali-E164G | 14 ± 1.1 | 1.6 ± 0.02 | 2.3 ± 0.2 | 4.2 |
| ML01[a] | 69 ± 5.6 | 1.5 ± 0.01 | 10 ± 0.9 | 20.4 |
| SenT190.08[a] | 32 ± 2.2 | 1.5 ± 0.02 | 4.7 ± 0.4 | 9.6 |

[a]Sequence from PlasmoDB

[b]Sequence from MalariaGen Pf7 dataset. Most common haplotype containing mutation of interest was expressed from the country identified.

**Table S16. Binding kinetics of Nb-Fc fusion proteins and bi-specific Nb-Fcs to PfPTRAMP-CSS**

| PfPTRAMP-CSS | | | | |
|---|---|---|---|---|
| **Nb-Fc** | **$K_D$ (nM)** | **$k_a$ (1/Ms) x $10^5$** | **$k_d$ (1/s) x $10^{-3}$** | **Fold $K_D$ (Average Nb-Fc/BsNb-Fc)[a]** |
| 1G2 | 11.5 ± 1.9 | 0.74 ± 0.21 | 0.82 ± 0.11 | - |
| 1G8 | 2.8 ± 0.5 | 7.8 ± 3.5 | 2.0 ± 0.63 | - |
| 2D11 | 23.1 ± 1.3 | 5.11 ± 0.22 | 11.9 ± 1.2 | - |
| 3E2 | 3.5 ± 1.3 | 5.02 ± 1.7 | 1.54 ± 0.06 | - |
| 3E2-1G2 | 0.001 ± 0.0 | 4.7 ± 1.4 | 0.0001 ± 0.0 | 7520 |
| 3E2-1G8 | 0.6 ± 0.2 | 0.59 ± 0.17 | 0.43 ± 0.13 | 5.4 |
| 3E2-2D11 | 0.2 ± 0.01 | 4.0 ± 0.59 | 0.073 ± 0.0161 | 7.5 |
| 1G8-2D11 | 0.2 ± 0.0 | 7.6 ± 0.24 | 0.13 ± 0.009 | 7.4 |
| 1G2-2D11 | 0.3 ± 0.0 | 3.94 ± 0.075 | 0.14 ± 0.023 | 5.1 |

[a]Compared to $K_D$ of nanobody only. For BsNb-Fcs the $K_D$ was compared to the average $K_D$ of both Nb-Fcs.

**Table S17. $EC_{50}$ (µg/mL) of Nb-Fc and BsNb-Fc growth inhibition activity in asexual blood-stage *P. falciparum* culture.**

| Nanobody-Fc | | |
|---|---|---|
| **Nb** | **$EC_{50}$ (µg/mL) (± 95% confidence interval)** | **Fold decrease in $EC_{50}$ (Lowest Nb-Fc/BsNb-Fc)** |
| 1G2-Fc | 556 (123 – 991) | - |
| 1G8-Fc | 64.4 (43.9 – 85.3) | - |
| 2D11-Fc | 540 (353 – 729) | - |
| 3E2-Fc | 558 (339 – 733) | - |
| **BsNb-Fc** | | |
| **Nb** | **$EC_{50}$ (µg/mL) (± 95% confidence interval)** | **Fold decrease in $EC_{50}$ (Lowest Nb-Fc/BsNb-Fc)** |
| 1G8-1G2 | 45.5 (28.95 - 88.1) | 1.4 |
| 1G2-3E2 | 73.7 (47.8 – 99.3) | 7.5 |
| 1G8-3E2 | 35.4 (16.2 – 73.7) | 1.8 |
| 2D11-3E2 | 18.1 (11.7 – 27.5) | 30 |
| 2D11-1G2 | 225 (127 – 315) | 2.4 |
| 1G8-2D11 | 44.4 (28.7 – 61.1) | 12 |
| R5.016 | 17.8 (16.1 – 21.4) | - |

*Compared with Nb-Fc with the lowest $EC_{50}$

**Table S18. Standard membrane feeding assays of Nb-Fcs and BsNb-Fcs tested at 100 µg/mL**

| Test Nb-Fc | Mean oocysts | No. of infected midguts/total midguts | %Transmission Reducing Activity (normalised to PBS) | Adjusted *p*-value[a] |
|---|---|---|---|---|
| **Experiment #1** | | | | |
| Untreated control (PBS) | 13.2 | 26/29 | 0 | - |
| WNb7-Fc (-) | 12.7 | 28/30 | 3.6 | >0.9999 (ns) |
| D2-Fc | 4.1 | 16/30 | 69.1 | 0.0001 |
| 1G2-Fc | 0.8 | 6/30 | 93.9 | <0.0001 |
| 1G8-Fc | 3.6 | 15/30 | 72.7 | <0.0001 |
| 1G12-Fc | 7.1 | 9/16 | 46.4 | 0.0412 |
| **Experiment #2** | | | | |
| Untreated control (PBS) | 28.4 | 29/30 | 0 | - |
| D2-Fc | 0.3 | 6/30 | 98.8 | <0.0001 |
| 1G2-Fc | 18.0 | 18/30 | 36.7 | 0.1601 (ns) |
| 1G8-Fc | 3.3 | 12/30 | 88.5 | <0.0001 |
| 1G12-Fc | 1.9 | 12/30 | 93.3 | <0.0001 |
| 3E2-Fc | 10.1 | 23/30 | 64.5 | 0.0028 |
| **Experiment #3** | | | | |
| Untreated control (PBS) | 28.2 | 31/31 | 0 | - |
| 2D11-Fc | 20.4 | 21/22 | 27.8 | >0.9999 (ns) |
| 3E2-Fc | 23.7 | 20/23 | 16.0 | >0.9999 (ns) |
| 3E2-2D11-Fc | 17.3 | 21/24 | 38.8 | 0.2633 (ns) |
| 1G2-2D11-Fc | 25.5 | 22/23 | 9.5 | >0.9999 (ns) |
| TB31F (+) | 0.1 | 3/24 | 99.6 | <0.0001 |
| **Experiment #4** | | | | |
| Untreated control (PBS) | 8.3 | 9/11 | 0 | - |
| 2D11-Fc | 7.7 | 17/22 | 6.6 | >0.9999 (ns) |
| 3E2-Fc | 11.4 | 21/23 | -37.7 | >0.9999 (ns) |
| TB31F (+) | 0.1 | 3/21 | 99.4 | 0.0016 |
| **Experiment #5** | | | | |
| Untreated control (PBS) | 25.7 | 26/27 | 0 | - |
| 3E2-2D11-Fc | 16.4 | 23/25 | 36.3 | 0.1599 (ns) |
| 1G2-2D11-Fc | 21.0 | 20/23 | 18.0 | 0.1352 (ns) |

[a] Kruskal-Wallis test compared to PBS untreated control number of oocysts

### Table S19. Sporozoite traversal assay

| Nanobody-Fc | Average % of dextran-FITC⁺ HC04 cells (n=2) | % *Pf* spz traversal normalised to WNb7-Fc | %Inhibition | % of dextran-FITC⁺ HC04 cells (n=1) | % *Pf* spz traversal normalised to WNb7-Fc | % Inhibition |
|---|---|---|---|---|---|---|
| Concentration | 100 µg/mL | | | 500 µg/mL | | |
| WNb36 (-) | 56.0 | 97.9 | 2.1 | 57.3 | 97.8 | 2.2 |
| WNb7 (-) | 57.2 | 100.0 | 0.0 | 58.5 | 100.0 | 0.0 |
| D2 | 56.7 | 99.1 | 0.9 | 57.6 | 98.5 | 1.5 |
| 1G2 | 57.4 | 100.3 | -0.3 | 56.9 | 97.3 | 2.7 |
| 1G8 | 56.5 | 98.7 | 1.3 | 57.6 | 98.3 | 1.7 |
| 2D11 | 57.0 | 99.6 | 0.4 | 58.9 | 100.6 | -0.6 |
| 3E2 | 59.0 | 103.1 | -3.1 | 58.6 | 100.1 | -0.1 |
| 3F4 | 59.2 | 103.5 | -3.5 | 56.9 | 97.2 | 2.8 |
| 3E2-1G8 | 55.6 | 97.1 | 2.9 | 57.0 | 97.4 | 2.6 |
| 3E2-2D11 | 52.5 | 91.8 | 8.2 | 50.8 | 86.8 | 13.2 |
| 1G2-2D11 | 53.7 | 93.9 | 6.1 | 55.1 | 94.2 | 5.8 |
| 3E2-1G2 | 52.4 | 91.5 | 8.5 | 55.8 | 95.3 | 4.7 |
| 1210 (+) | 4.8 | 8.3 | 91.7 | 6.2 | 10.6 | 89.4 |

+ positive control monoclonal antibody

### Table S20. Sporozoite hepatocyte invasion assay with anti-PTRAMP-CSS Nb-Fcs.

| Donor AY22 | | | | Donor AY17 | | |
|---|---|---|---|---|---|---|
| Nanobody-Fc or Antibody | Average %parasite infected hepatocytes | %Infected hepatocytes (normalised to WNb7-Fc) | %Sporozoite invasion inhibition | Average %parasite infected hepatocytes | %Infected hepatocytes (normalised to WNb7-Fc) | %Sporozoite invasion inhibition |
| 1G8 | 0.4 | 38.4 | 61.6 | 0.9 | 48.4 | 51.6 |
| 2D11 | 0.9 | 82.8 | 17.2 | 1.7 | 96.9 | 3.1 |
| 3E2 | 0.7 | 65.8 | 34.2 | 1.3 | 74.9 | 25.1 |
| 1G2 | 0.7 | 62.6 | 37.4 | 1.2 | 64.8 | 35.2 |
| D2 | 0.9 | 80.9 | 19.1 | 1.9 | 106.9 | -6.9 |
| 3F4 | 1.0 | 89.6 | 10.4 | 1.8 | 100.7 | -0.7 |
| 2D11-3E2 | 0.5 | 50.1 | 49.9 | 0.9 | 50.4 | 49.6 |
| 3E2-1G2 | 0.6 | 56.3 | 43.7 | 1.1 | 61.2 | 38.8 |
| 2D11-1G2 | 0.5 | 45.7 | 54.3 | 1.0 | 54.1 | 45.9 |
| 1G8-3E2 | 0.4 | 39.5 | 60.5 | 0.9 | 51.9 | 48.1 |
| HIHS (-) | 0.8 | 72.4 | 27.6 | 1.9 | 107.8 | -7.8 |
| 1210 (+) | 0.0 | 1.3 | 98.7 | 0.0 | 2.2 | 97.8 |
| WNb7 (-) | 1.1 | 100.0 | 0.0 | 1.8 | 100.0 | 0.0 |
| WNb36 (-) | 1.1 | 101.0 | -1.0 | 1.8 | 101.7 | -1.7 |

+ denotes positive control antibody and – denotes negative control Nb-Fcs
